# Patient-derived organoids to study glycosylation dynamics during gastric disease

**DOI:** 10.1101/2025.02.21.639100

**Authors:** Liliana Santos-Ferreira, Henrique O. Duarte, Eva Moia, Ana F. Costa, Álvaro M. Martins, Isabel Faria-Ramos, Rita Matos, Rita Barros, Marta Silva, Sofia Pedrosa, Diana A. Batista, Joana Gomes, Thomas Borén, Fabiana Sousa, Frederica Casanova-Gonçalves, José Barbosa, Ana Magalhães, Catarina Gomes, Hugo Santos-Sousa, Sina Bartfeld, Fátima Carneiro, Celso A. Reis, Filipe Pinto

## Abstract

**BACKGROUND AND AIMS:** Aberrant cellular glycosylation remains a key event that accompanies and actively sustains gastric neoplastic transformation. Patient-derived organoids (PDOs) have recently emerged as a promising *ex vivo* model to study human gastric disorders. Since the PDOs glycosylation landscape remains unknown, this study aims to evaluate PDOs as potential *avatars* of *in vivo* tissue glycosylation profiles in the gastric context.

**METHODS:** Fresh gastric mucosa samples derived from non-tumoral obese patients (n=11), adjacent tumor mucosa samples (n=29), and tumor tissue samples derived from gastric cancer (GC) patients (n=30) were used to establish a biobank of gastric PDOs (n=56). The *N*- and *O-*glycophenotypes of normal, adjacent, and tumor PDOs and respective *in vivo* tissues were thoroughly characterized by immunostaining. Additionally, a comparative glycan analysis was performed over time, upon PDO biobanking and xenografting in mice. The binding of two *Helicobacter pylori* (*H. pylori*) isogenic strains with distinct glycan-binding affinities was assessed in parental gastric mucosa tissues and compared with the respective PDOs before and after modulation of their glycan landscape.

**RESULTS:** Our results show that PDOs mimic different phenotypes of the carcinogenic cascade and recapitulate parental gastric tissues’ glycosylation profile. Tumor PDOs recapitulate the inter- and intra-heterogeneity features observed in GC, which is maintained over time, upon biobanking and xenografting. We demonstrated that the expression of type I and type II Lewis antigens is dynamically controlled by PDOs differentiation status, which results in differential binding to *H. pylori* strains displaying distinct glycan-binding adhesins, mirroring the gastric epithelium tissue interactions.

**CONCLUSIONS:** This study established PDOs as invaluable *ex vivo* tools to study the complex glycan dynamics in both gastric physiological and pathological settings.

## INTRODUCTION

Aberrant glycosylation plays a crucial role in the onset and progression of gastric malignancy, as highlighted by us and others^1–4^. Indeed, *Helicobacter pylori* (*H. pylori*) infection, one of the major risk factors associated with the development of gastric cancer (GC)^5, 6^, is mediated by the binding of the adhesin-equipped bacteria to specific glycan moieties, whose distribution is highly regulated across the gastric epithelium^7^. The ABO/Lewis b (Le^b^) histo-blood group antigens, and their major molecular carrier mucin MUC5AC, act as an anchoring epitope for the bacterial blood group antigen binding adhesin (BabA)^8–10^. Persistent *H. pylori* infection triggers chronic gastric inflammation, characterized by the *de novo* expression of sialylated glycans, particularly sialyl-Lewis A (sLe^a^) and sialyl-Lewis X (sLe^x^)^11–13^. These inflammation-induced glycan antigens serve as binding epitopes to the bacteria’s sialic acid-binding adhesin (SabA)^11^, ultimately functioning as additional anchoring points of *H. pylori* to the inflamed gastric epithelium^11, 13–15^. Gastric carcinogenesis evolves from chronic gastritis into pre-malignant conditions, such as intestinal metaplasia (IM), which is accompanied by the anomalous *de novo* expression of mucin MUC2 carrying the truncated *O-*glycan sialyl-Tn (sTn)^16–18^. In malignant conditions, other glycans are aberrantly expressed, which includes the overexpression of short *O*-glycans, the increase of sialylated and fucosylated structures, and an increase in the branching patterns of *N*-linked glycans^2^. These glycan alterations have been shown to affect tumor aggressiveness, therapeutic response, and patient clinical outcome ^3, 19–26^. Taking into consideration the impact of glycans in GC, there is a need for advanced disease models to study the complexity and dynamics of glycosylation throughout gastric carcinogenesis.

In recent years, patient-derived organoids (PDOs) have emerged as promising *ex vivo* models to study cellular and molecular processes underlying human gastric disorders^27, 28^. Particularly, PDOs have been a valuable tool for studying *H. pylori* infection in the context of gastric pathology^29–31^ and for understanding GC response to drug treatment^32–34^. However, despite the importance of glycosylation and the potential for PDOs to offer insights into gastric disorders, the role of this dynamic cellular process within gastric PDOs remains poorly explored. In this study, we address this knowledge gap by studying the glycosylation landscape in human gastric PDOs derived from normal mucosa, adjacent tumor mucosa, and tumor tissues, to provide novel insights into gastric carcinogenesis as well as on tumor progression.

## MATERIAL AND METHODS

### Human tissue samples

Fresh normal gastric mucosa was obtained from individuals undergoing gastric sleeve resection. Tumor, and corresponding adjacent mucosa were obtained from patients undergoing surgical gastrectomy/tumor resection at Unidade Local de Saúde (ULS) de São João in Porto, Portugal (ethical approved reference number CES 223/2021). A total of 70 human tissue samples were collected in this study. The cohort comprises 11 gastric mucosa samples from non-tumoral obese patient donors (N) and 59 samples from patients diagnosed with GC, including 29 samples from gastric mucosa adjacent to the tumor (Adj) and 30 samples from tumor tissues (T). Patient clinicopathological information and additional details of material and methods are described in **Supplementary Table 1, Supplementary Table 2** and **Supplementary Methods**.

### Mice

C57BL/6 *wild-type*, *Gcnt1^−/−^* and nude CBA mice were bred and housed at i3S animal facility, Portugal. *Gcnt1^−/−^* were obtained from the Consortium for Functional Glycomics (CFG)^35^. Additional information is provided in the **Supplementary Methods**.

### Gastric mice and human patient-derived organoids, and xenograft generation

Normal gastric mice organoids (MDO) and normal human PDOs were performed as described previously^27^. The protocol to generate tumor organoids was adapted from a previously described protocol from Wallascheck, *et al.*^36^, and is described in the **Supplementary Methods**. All materials and methods to evaluate organoid differentiation are described in **Supplementary Table 3**, **Supplementary Table 4** and **Supplementary Methods.** Eleven gastric tumor PDOs (T-PDOs) with different histologic phenotypes (solid, glandular, non-cohesive and mixed pattern) were used to generate patient-derived tumor organoids xenografts (PDOX) as described previously^37^. See **Supplementary Table 1** and **Supplementary Table 2** for clinicopathological data and histologic characterization of PDOs. Detailed methods are described in **Supplementary Methods**.

### Helicobacter pylori strains and binding assays

The *H. pylori* strains 17875/Le^b^ and 17875babA1::kan babA2::cam (17875 DM) were obtained from the Department of Medical Biochemistry and Biophysics, Umeå University, Sweden^11^. These *H. pylori* strains were used to functionally evaluate the impact of PDOs glycophenotypes on *H. pylori* binding. Detailed material and methods are described in **Supplementary Methods**.

### Glycan profiling of patient-derived organoids

PDOs and their respective tissue samples were analyzed by immunofluorescence, immunohistochemistry, and lectin-based histochemistry. Detailed information about antibodies, lectins and specific staining protocols is described in **Supplementary Table 4** and **Supplementary Methods**.

## RESULTS

### Establishment of patient-derived organoids from normal, non-tumoral adjacent mucosa and tumor gastric specimens

PDOs have been described as an *ex vivo* model to study human gastric mucosa development and associated diseases^27–34, 36, 38^. However, the understanding of the role of glycosylation within this 3D model remains limited, precluding its full exploitation in cancer research and in pre-clinical settings. To address this gap, we generated a total of 56 gastric PDO cultures, including 11 PDOs from normal gastric mucosa (N-PDO) derived from obese patients, with a success rate of 100%; 26 PDOs from adjacent gastric mucosa (Adj-PDO) and 19 tumor PDOs (T-PDO) derived from patients diagnosed with GC with a success rate of 89% and 63 %, respectively (**Figure 1A)**. From the Adj-PDOs established, six of them are “sibling” organoids derived from two different adjacent tissue specimens from three patients (patient Adj11, Adj16, and Adj22; **Supplementary Table 1**), as well as twelve “sibling” T-PDOs from two different tumoral regions from six tumor specimens (patient T1, T12, T15, T20, T21, and T22; **Supplementary Table 2)**. A comprehensive overview of the established biobank currently available in our laboratory is provided in **Supplementary Figure 1**. The clinicopathological information from the patients used in this study is provided in **Supplementary Table 1** and **Supplementary Table 2**.

**Figure 1.**
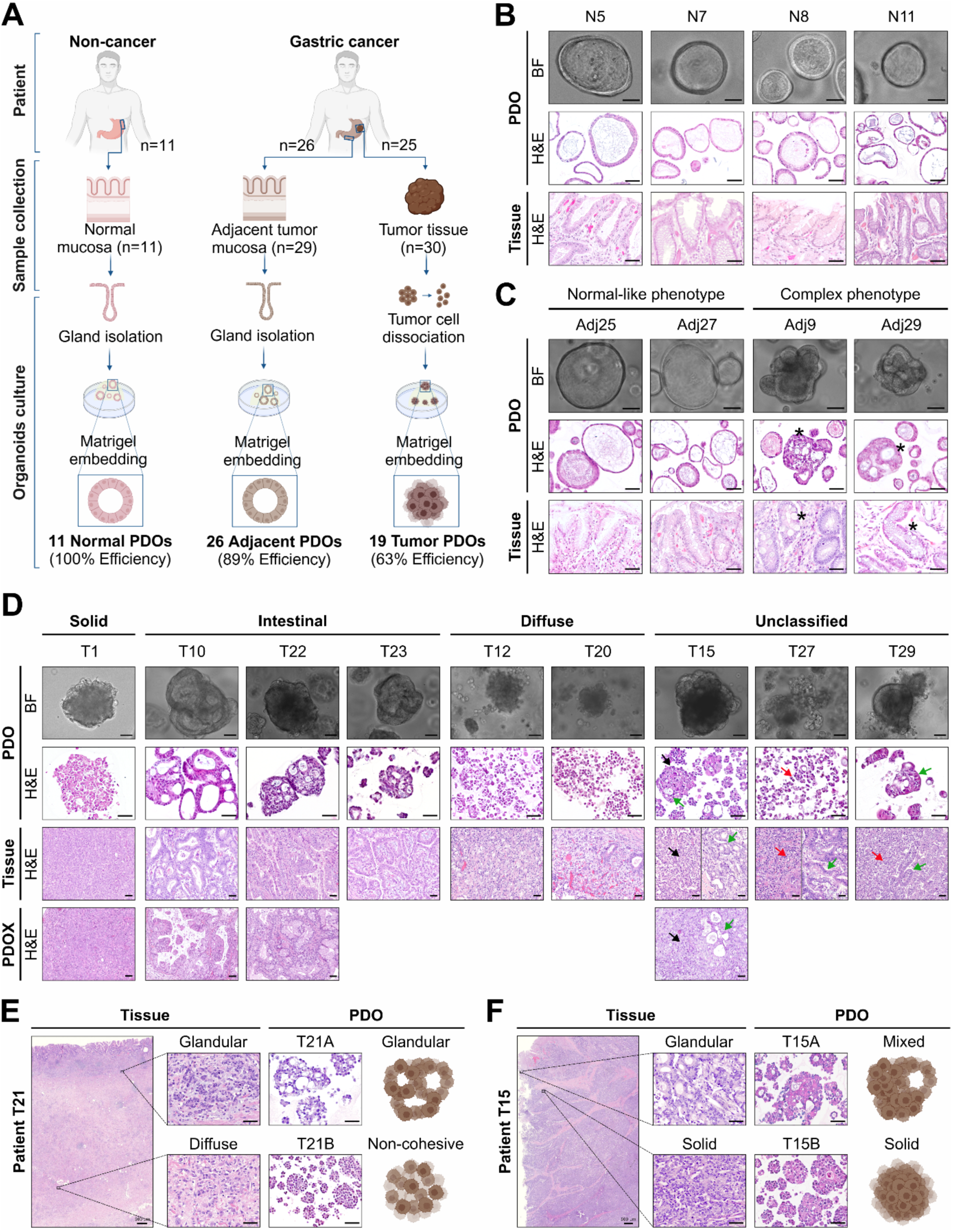
Normal, adjacent and tumor gastric patient-derived organoid biobank. **A)** Graphical representation of patient sample collection and patient-derived organoids (PDOs) biobank approach. **B)** Representative brightfield (BF) and hematoxylin-eosin staining (H&E) images of PDOs derived from normal gastric mucosa (N). **C)** Representative BF and H&E images of PDOs derived from non-tumoral gastric mucosa adjacent to the tumor (Adj). Two main phenotypes were observed in the Adj-PDOs: normal-like and complex structures akin to those found in parental tissues. * on Adj-PDOs indicates premalignant phenotypes such as IM/dysplasia-like morphologies, recapitulating the parental tissues; * on tissues indicates IM on gastric glands. **D)** Representative images of tumor (T) PDOs established in the present work from different gastric cancer subtypes classification (solid, intestinal, diffuse, and unclassified subtypes). T-PDOs reflect the histologic features of original tissues maintained upon xenograft PDOs in mice models (PDOX). Arrows indicate the different histologic components observed in PDOs and tissues: the black arrow indicates the solid component; the green arrows indicate the glandular component and the red arrows indicate the non-cohesive component. **E-F)** “Sibling” T-PDOs derived from different tumor tissues of patient T21 (**E**) and patient T15 (**F**) diagnosed with unclassified subtype (mixed) of gastric cancer. Scale bar represents 50 µm.

The isolated glands from gastric mucosa (N-PDOs and Adj-PDOs) give rise to cystic structures characterized by a single layer of epithelial cells surrounding the central lumen (**Figure 1B-C** and **Supplementary Figure 1**), as described^27, 36^. Interestingly, distinct morphological phenotypes within the same culture were also observed, especially among the Adj-PDOs. These PDOs exhibited an altered and complex re-arrangement with focal irregularity and cribriform-like structures with some lacking a well-defined lumen (**Figure 1C, Supplementary Figure 1** and **Supplementary Figure 2A**). Specifically, 20% N-PDOs (2 out of 10) and 60% of Adj-PDOs (15 out of 25) depicted a complex phenotype representing less than 20% of the PDO population (**Supplementary Table 1**).

The T-PDOs exhibited a heterogeneous, and non-organized cellular structure while preserving the histomorphology subtype of the original gastric adenocarcinoma tissues (**Figure 1D, Supplementary Table 2**), as described^32, 36^. Based on their morphology and histological architecture, T-PDOs were classified into solid, glandular, non-cohesive, and mixed (glandular/solid and non-cohesive/solid) subtypes, recapitulating the original tissue histology, which was further confirmed upon T-PDO xenografting in mice (PDOX) (**Figure 1D**, **Supplementary Figure 2B**, and **Supplementary Table 2**).

Interestingly, “sibling” T-PDOs derived from different tumor regions of the same patient diagnosed with mixed GC subtype exhibited distinct morphologies. Specifically, T-PDO 21A has a glandular morphology while T-PDO 21B presents a non-cohesive and solid pattern, which is aligned with the original mixed histological subtype displaying diffuse and glandular components (**Figure 1E**). This was also observed in patient T15, where T-PDO 15A exhibits a predominantly glandular component, as opposed to the major solid component of “sibling” T-PDO 15B (**Figure 1F**). On the other hand, the “sibling” T-PDOs derived from samples with well-defined histological subtype - solid (T-PDO 1 and 1.1), intestinal (T-PDO 22 A and B) and diffuse (T-PDO 12 A and B; T-PDO 20A and B) - show histological similarity between “sibling” T-PDOs (**Supplementary Figure 2C** and **Supplementary Table 2**). Taken together, our PDO biobank mimics the histological and morphological features, as well as the heterogeneity of the derived tissues, ranging from normal to tumor-adjacent and tumor tissue.

### Adjacent patient-derived gastric organoids display features of gastric pre-malignant conditions

Non-tumoral PDOs (N- and Adj-PDOs) showed a consistent expression of gastric tissue molecular markers, such as mucins MUC5AC and MUC6, pepsinogen C (PGC), and somatostatin (SST) suggesting that these PDOs follow the normal epithelium gastric cell differentiation **(Figure 2A** and **Supplementary Figure 3A-C**). However, complex morphologies depicting vacuole and cribriform-like structures, especially in the Adj-PDOs derived from tumor-adjacent mucosa (**Figure 1C** and **Supplementary Figure 2A**), resembling histological features of gastric pre-malignant conditions such as intestinal metaplasia (IM) or dysplasia^39^, were observed. This prompted us to investigate whether these PDOs could recapitulate these pre-malignant conditions *ex vivo*.

**Figure 2.**
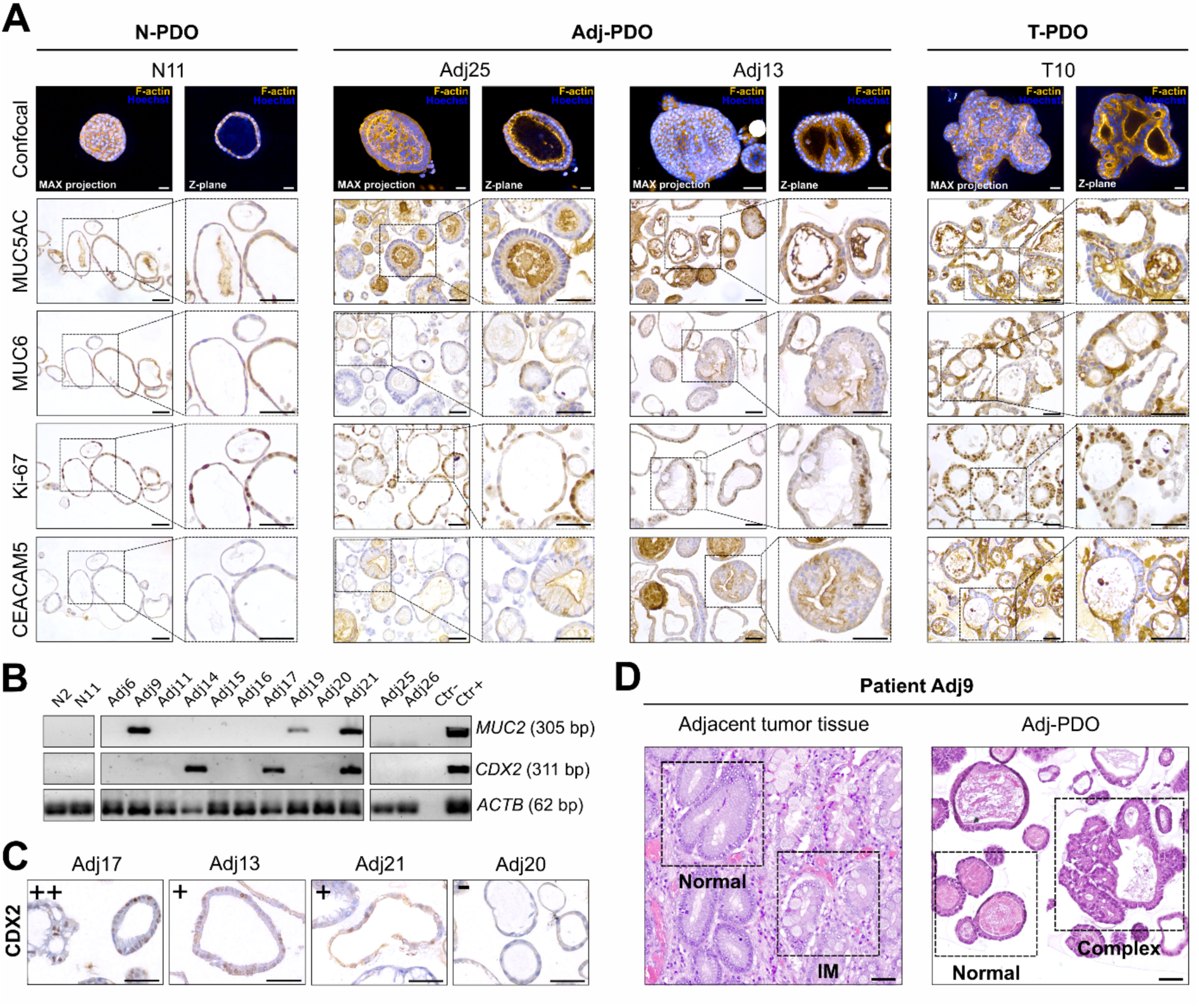
Non-tumoral gastric organoids recapitulate distinct features of gastric carcinogenesis. **A)** Representative confocal immunofluorescence and immunohistochemistry images of PDOs showing distinct gastric PDO phenotypes. N-PDOs (N11) express classical staining for mucin MUC5AC and MUC6, while Adj-PDOs (Adj25 and Adj13) show increased levels of proliferation (Ki-67) and of the tumoral marker CEACAM5, which are maintained in the T-PDOs (T10). Max projection and Z-plane indicate the maximum intensity projection and the individual section of the confocal images, respectively. **B)** mRNA expression of pre-malignant markers of intestinal metaplasia (*MUC2* and *CDX2*) in two N-PDO and twelve Adj-PDOs **C)** Representative CDX2 immunostaining in four Adj-PDOs, demonstrating the nuclear expression of CDX2 in three Adj-PDOs with differential staining levels (Adj17, Adj13 and Adj21) and one Adj-PDO (Adj20) negative for CDX2. Human tissue with IM lesions was used as a positive control (CTR+). **D)** H&E staining of the parental adjacent gastric mucosa and respective derived Adj-PDOs from patient Adj9. The same tissue fragment can show both normal and premalignant lesions (as intestinal metaplasia), which is reflected in the two different Adj-PDOs (normal *vs* complex phenotypes) that originated from that tissue. Scale bar represents 50 µm.

For that, the expression of the classical IM molecular markers of intestinal transdifferentiation *CDX2* and/or mucin *MUC2,* the proliferative markers Ki-67 and the cancer-associated marker CEACAM5 were assessed in the PDOs (**Figure 2A-C)**. Interestingly, 46% of the tested Adj-PDOs (n=6/13) express the IM molecular markers *CDX2* and/or mucin *MUC2* (**Figure 2B -C**). These data support the presence of a PDO IM-like subpopulation. Moreover, we found that both Adj-PDOs and T-PDOs showed high staining of both Ki-67 and CEACAM5 markers compared with N-PDOs (**Figure 2A** and **Supplementary Figure 3A**). These molecular data, combined with the observed heterogeneous morphological phenotypes, indicate that, from the same organoid culture, PDOs with a normal-like and/or with a pre-malignant phenotype can be simultaneously developed (**Figure 2D** and **Supplementary Table 1**). Such heterogeneous populations of organoids may stem from the gastric lesion heterogeneity found in most patients, and from which gastric PDOs are established. Indeed, several collected adjacent gastric mucosa simultaneously display healthy glands and regions depicting pre-malignant morphologies such as IM and dysplasia (**Figure 2D** and **Supplementary Table 1**), which may originate PDOs with both morphological and molecular phenotypes.

### Patient-derived organoids recapitulate the in vivo glycophenotype of normal and pre-malignant gastric epithelial glands

To evaluate how closely non-tumoral PDOs resemble the glycan profile of parental epithelium tissues, we performed immunostaining analysis in N- and Adj-PDOs, and the corresponding parental tissues for key glycan antigens found in the gastric epithelium, including type I Lewis antigens, type II Lewis antigens, fucosylated structures, sialylated structures, branched *N*-glycans, bisecting *N*-glycans, and short truncated *O*-glycans structures (**Supplementary Table 4)**.

The described differences between tumor adjacent and normal mucosa tissues, as increased levels of Lewis a (Le^a^), sialylated lewis (sLe^a^ and sLe^x^) and truncated O-glycans (T and sTn)^40–42^ were maintained in the Adj-PDOs compared with N-PDOs (**Figure 3A**). No significant differences were observed for other glycans when comparing normal with adjacent samples (**Figure 3A** and **Supplementary Figure 4A**). However, when comparing the glycan levels between tissues and PDOs, it was observed an enrichment of sialylated Lewis antigens (sLe^a^ and sLe^x^) in Adj-PDOs, and increased Le^a^ levels in N-PDOs compared with parental tissues (**Figure 3A)**. Such differences may be a consequence of the tissue fragment from which specific PDOs were generated, differential organoid maturation, selective culture conditions, or selective biological advantages promoted by these specific glycans to cells. Overall, the observed glycan staining patterns consistently showed that N- and Adj-PDOs mimic the corresponding parental tissues glycoprofile (**Figure 3A** and **Supplementary Figure 4A**). The gastric epithelial PDO glycoprofile was also maintained upon genetic manipulation, as observed in mice-derived organoids (MDOs) from the gastric mucosa that either expressed or lacked the Gcnt1 glycosyltransferase (**Supplementary Figure 4B-D**). Knockout (KO) mice models for *Gcnt1* lack core 2 extension resulting in the accumulation of core 1 glycan structures such as T and Tn antigens^35^, which were maintained in the respective KO MDOs (**Supplementary Figure 4B-C**). No major differences were observed for other glycan epitopes, except for sLe^x^ and Le^y^, which showed overrepresented staining in both KO gastric tissue and KO MDOs (**Supplementary Figure 4D**). Taking together, these data validate PDOs as a reliable model for the recapitulation of gastric tissue glycophenotypes.

**Figure 3.**
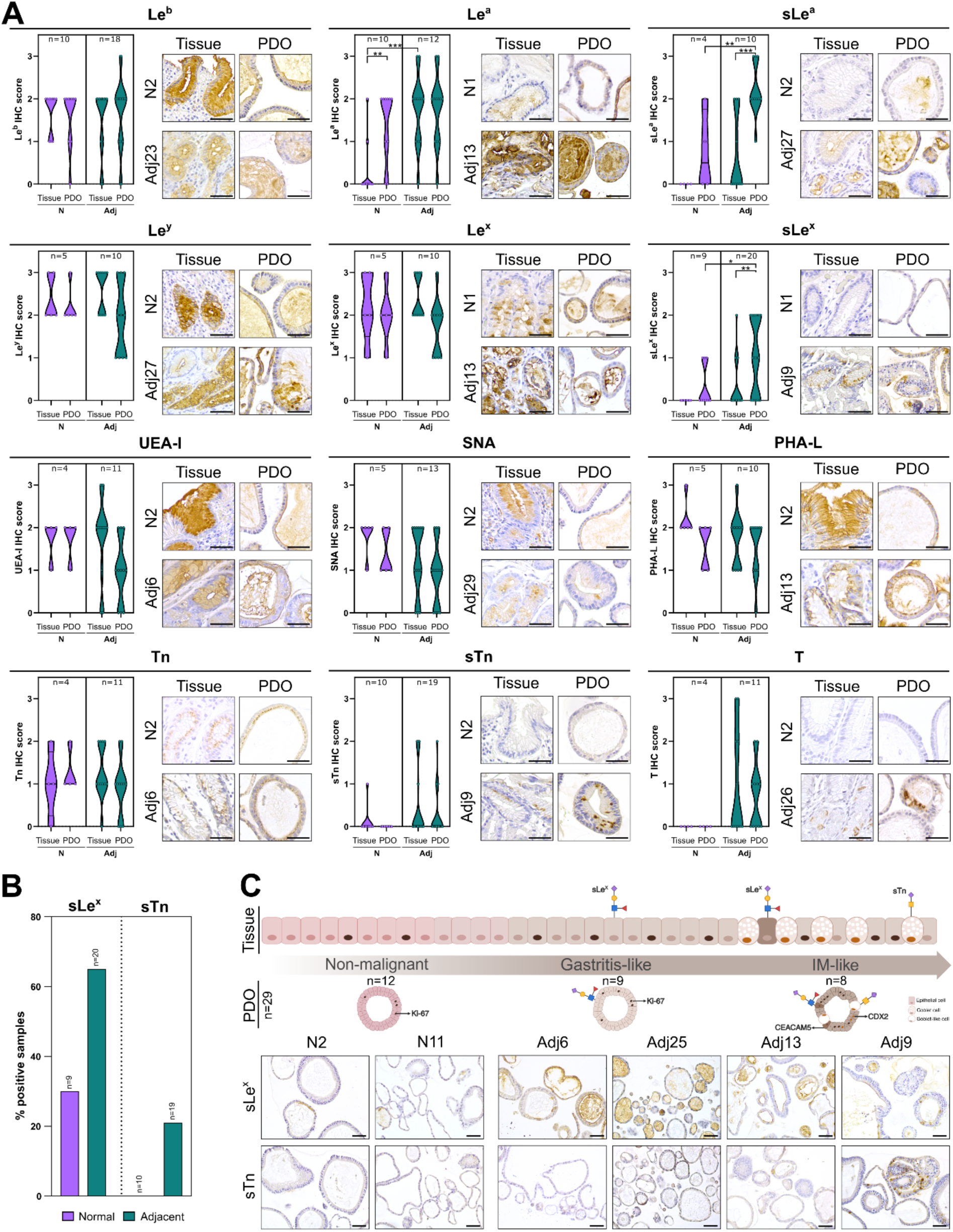
The glycophenotype of normal and adjacent gastric mucosa tissues can be recapitulated by respective normal and adjacent-derived organoids. **A)** Graphical quantification of glycan immunostaining and representative images of the immunostaining are represented for type I Lewis antigens (Le^a^, Le^b^, sLe^a^), type II Lewis antigens (Le^x^, Le^y^, sLe^x^), terminal fucosylated structures (*Ulex europaeus agglutinin-I* - UEA-I), α2,6 sialylated structures (*Sambucus nigra* lectin - SNA), branched *N*-glycans (*Phaseolus vulgaris* leucoagglutinin - PHA-L), and short *O*-glycans (Tn, sTn, and T) found in normal (N), adjacent (Adj) PDOs and matched tissues. *p value*<0.05 was considered statistically significant (unpaired t-test; * indicates a *p value*<0.05; ** indicates a *p value*<0.005, and *** indicates a *p value*<0.001). **B)** Percentage of N- and Adj-PDOs showing positive staining for sLex and sTn. **C)** Schematic representation of premalignant glycophenotype found in gastric carcinogenesis and PDOs. Representation of the sLe^x^ and the truncated *O*-glycan sTn glycan staining in PDOs are represented below. Scale bar corresponds to 50 µm.

The canonical gastric carcinogenic cascade is characterized by the *de novo* expression of sialylated glycans, such as: sialyl Lewis X (sLe^x^), whose biosynthesis by gastric epithelial cells is triggered by persistent *H. pylori* infection^11^; and sialyl Tn (sTn), whose expression is induced in regions of the mucosa undergoing IM^16–18^. We thus evaluated whether the PDOs could recapitulate these specific glycophenotypes. sLe^x^ showed to be positive in 30.0% of the N-PDOs (3/10) and 65.0% of the Adj-PDOs (13/20). Moreover, none of the N-PDOs expressed sTn (0.0%, 0/10), while 21.0% of the Adj-PDOs (4/19) expressed sTn (**Figure 3B)**. These results indicate that gastric PDOs can reflect different glyco-stages of the gastric carcinogenesis. As such, the biobank was categorized into normal (negative for both sLe^x^ and sTn, n=12, gastritis-like (sLe^x^-positive and sTn-negative, n=9), IM-like (sTn-positive and/or CDX2/MUC2-positive, n=8), and tumoral (n=19) (**Figure 3C**). The morphological and phenotypic similarities between specific organoids and gastric pre-malignant conditions show that organoids are a valuable *ex vivo* model to explore gastric carcinogenesis.

### Lewis antigens glycan profiles are dependent on patient-derived organoids differentiation and influence glycan-dependent Helicobacter pylori binding

Considering the observed enrichment of sialylated type I and type II Lewis antigens in PDOs compared to paired tissues, and knowing that these antigens are expressed in distinct gastric cell populations (foveolar cells - Lewis type I; and mucous gland cells - Lewis type II)^43^, we hypothesized that such differences might be attributed to PDO differentiation status. Phenotypical differentiation was induced in three PDOs (N-PDO 2, N-PDO 11 and Adj-PDO 9) derived from tissues showing neglectable/ low expression levels of these glycans (**Supplementary Figure 5A**). PDO differentiation was performed by altering the WNT3A concentration in the culture media^27^, which stimulates the differentiation of either foveolar pit-type organoids or mixed gland-type organoids (**Figure 4A** and **Supplementary Figure 5B**). PDOs differentiated into foveolar pit-type organoids (MUC5AC-positive and MUC6-negative) depicted upregulated levels of type I Lewis antigens (Le^b^ for all PDOs and sLe^a^ for Adj9), and loss of expression of type II Lewis antigens (Le^x^ and sLe^x^ for all PDOs) when compared with the mixed-type organoids (MUC5AC- and MUC6-positive) (**Figure 4A** and **Supplementary Figure 5B**). In PDOs with mixed populations, both type I and type II Lewis antigens are maintained in around 40-50 % of organoids resembling the mucosal heterogeneity of cell types expressing MUC5AC and MUC6 (**Figure 4A** and **Supplementary Figure 5B**). Interestingly, the small undifferentiated PDOs (4 days of growth) show to be highly positive for the expression of sialylated Lewis antigens (**Supplementary Figure 5C**). This data found in undifferentiated PDOs indicates that sLe^x^ and sLe^a^ may be involved in stem-like PDOs development.

**Figure 4.**
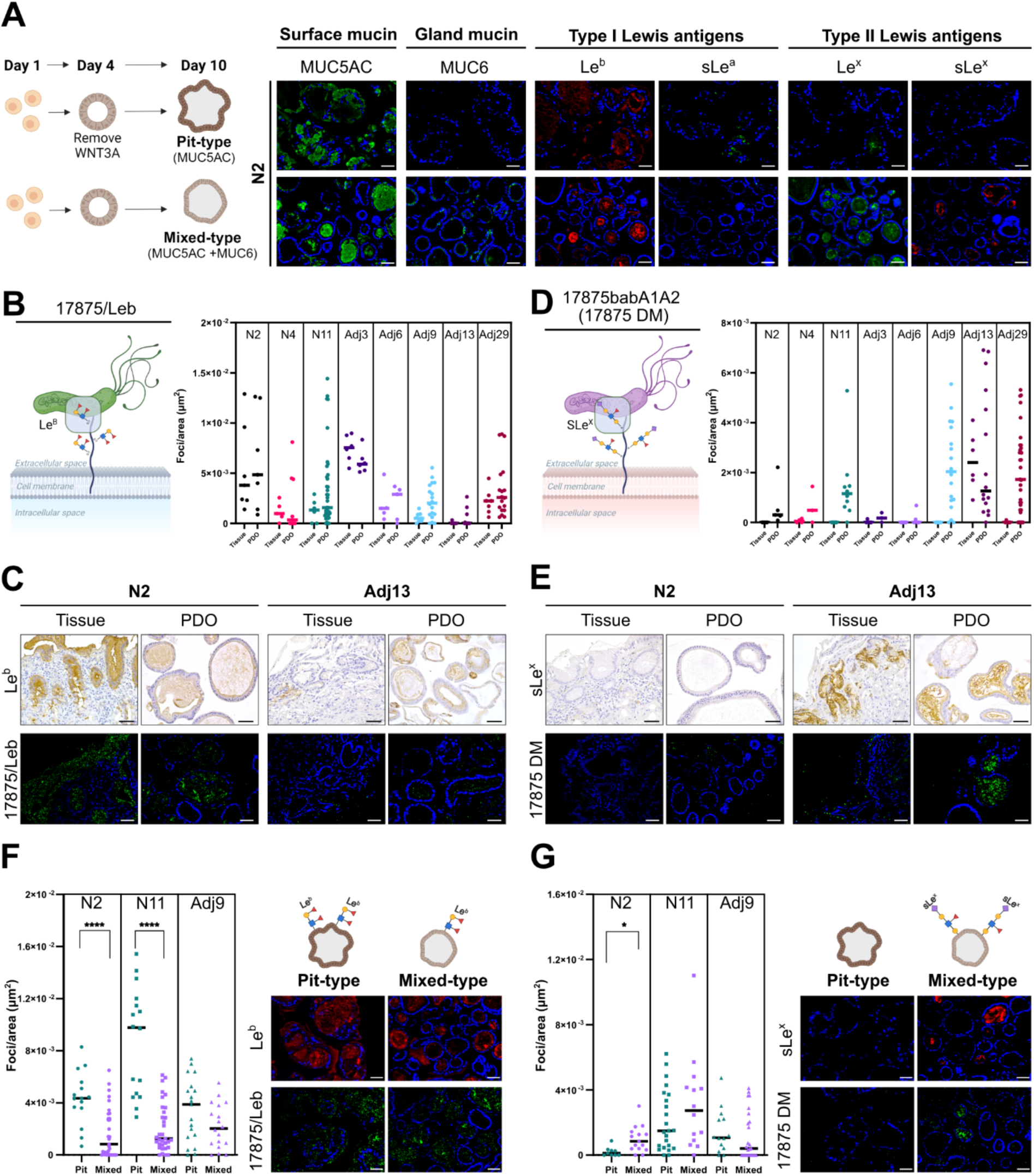
Type I and type II Lewis antigens expression is dependent on gastric organoid differentiation status and impact *H. pylori* binding. **A)** On left is a schematic representation of the PDO differentiation assay into foveolar pit-type (no WNT3A) and gland mixed-type (with WNT3A). Illustrative staining of surface mucin MUC5AC, gland mucin MUC6, type I and II Lewis in N-PDO 2. **B)** Schematic representation of the 17875/Leb *H. pylori* strain binding to the Lewis b (Le^b^) antigen. On the right is the quantification of 17875/Leb *H. pylori* binding per area (µm^2^) in the epithelium of normal and adjacent tissues and their respective PDOs. **C)** Representative images of 17875/Leb *H. pylori* binding in one Le^b^-high (N2) tissue and one Le^b^-low (Adj13) tissue and respective PDOs. **D)** Schematic representation of the double mutant 17875babA1/A2 (17875 DM) *H. pylori* strain binding to sialylated Lewis antigens, such as sLe^x^. On the right is the quantification of 17875 DM binding per area (µm^2^) in the epithelium of normal and adjacent tissues and their respective PDOs. **E)** Representative images of DM *H. pylori* binding in one sLe^x^-negative (N2) tissue and one sLe^x^-positive (Adj13) tissue and respective PDOs. **F)** Quantification of 17875/Leb strain binding per area (µm^2^) in differentiated PDOs and respective immunofluorescence staining of Le^b^ antigen. **G)** Quantification of DM strain binding per area (µm^2^) in differentiated PDOs, and respective immunofluorescence of sLe^x^ antigen. Scale bar corresponds to 50 µm. *p value*<0.05 was considered statistically significant (unpaired t-test; * indicates a *p value*<0.05; and **** a *p value*<0.0001).

It is known that *H. pylori* binding and tropism across the gastric mucosa are determined by the Lewis antigens glycosylation patterns of the gastric cells^7^. As such, we evaluated whether PDO Lewis antigen profile impacts *H. pylori* binding capacity. To investigate this, a binding assay was conducted using two isogenic *H. pylori* mutants on N- and Adj-PDOs, and the corresponding tissues. The BabA-expressing 17875/Leb strain binds to Le^b^ but does not bind to sialylated antigens, whereas the SabA-expressing 17875 DM strain binds to sialylated Lewis antigens, such as sLe^x^ and sLe^a11^. We observed that 17875/Leb binding to both healthy and adjacent tissues is retained in N- and Adj-PDOs, respectively, according to Le^b^ antigen expression (**Figure 4B-C** and **Supplementary Figure 6A**). The 17875 DM strain demonstrated higher binding activity to sLe^x^-positive Adj13 tissue as well as to its respective Adj-PDO (Adj-PDO13). No significant binding was observed in the negative and weakly positive sLe^x^ tissues; however, the 17875 DM strain still binds to the majority of corresponding PDOs (**Figure 4D-E** and **Supplementary Figure 6B**). This binding is mainly due to the sLe^x^ and sLe^a^ expression in more undifferentiated PDOs (**Supplementary Figure 5C**). Interestingly, it was noted that 17875/Leb binds more to both normal tissues and their respective N-PDOs compared to adjacent samples, whereas 17875 DM binds more effectively to adjacent tissues and their respective Adj-PDOs while binding less to normal tissues and PDOs (**Figure 4B and 4D**). This demonstrates that *H. pylori* tropism from normal tissue toward adjacent tumor tissues is maintained in PDOs.

Then, *H. pylori* binding assays were also performed in the PDOs with controlled differentiation status and specific type I or type II Lewis antigen glycoprofiles, as described above (**Figure 4A** and **Supplementary Figure 5B**). Upon organoid differentiation, we observed that the binding of the 17875/Leb strain was higher in all PDOs differentiated into foveolar pit-type (enriched in Le^b^-positive organoids), compared with the mixed-type organoids (heterogeneous expression of Le^b^) (**Figure 4F** and **Supplementary Figure 6C**). In contrast, increased 17875 DM strain binding was observed in enriched-sLe^x^ mixed-type N-PDOs (N2 and N11), compared with pit-type PDOs (negative for sLe^x^ and sLe^a^) (**Figure 4G**, **Supplementary Figure 6D**). The pre-malignant Adj-PDO 9 showed no significant differences in 17875 DM binding (**Figure 4G**). This observation is most probably due to the high sLe^a^ expression levels observed in both mixed- and pit-type PDOs (**Supplementary Figure 5B**), which also serves as an anchoring epitope for the 17875 DM strain.

Together the alignment in glycan expression, PDO differentiation, and *H. pylori* binding highlights the PDOs fidelity in recapitulating the complex and dynamic glycophenotype of normal and pre-malignant gastric epithelia.

### Specific gastric cancer glycan heterogeneity is preserved in tumoral patient-derived organoids

To assess whether tumor PDOs can be used as a robust model for studying glycosylation in cancer research, we then evaluated their ability to retain tumor tissue glycoprofile for the same glycan antigens described above. Our findings indicate that tumor PDOs effectively preserve the glycan expression patterns observed in the corresponding tumor tissues independently of the GC subtype (**Figure 5** and **Supplementary Figure 7A**). Overall, intestinal GC tissues showed increased levels of truncated *O*-glycans (Tn, sTn and T) and sialylated Lewis antigens (sLe^x^ and sLe^a^), while the diffuse GC subtype tumors showed a prevalence of sLe^a^, α2,6-sialylated Lac(di)NAc (SNA) and, β1,6 branched *N*-glycans (PHA-L). The unclassified mixed subtype tumors present a heterogeneous glycan expression pattern for all glycan epitopes studied, which reflects the mixed intestinal and diffuse nature of these tumors. On the other hand, the tumors with solid pattern (undifferentiated tumors) are highly positive for α2,6-sialylated Lac(di)NAc (SNA) and for the truncated *O*-glycan T, but with focal expression of all other glycans studied. The generated T-PDOs follow the same pattern of glycan expression mimicking the GC subtypes from where they originated. Importantly, T-PDOs also preserve the different tumor cell clonal subpopulations with distinct glycophenotypes frequently observed within the tumor tissues (**Figure 5** and **Supplementary Figure 7A**). Moreover, when comparing the glycoprofiles of tumor tissues and PDOs with normal and adjacent samples, we observed a progressive increase in sialylated Lewis antigens (sLe^a^ and sLe^x^), truncated *O-*glycans (sTn and T), and Le^a^, along with a progressive decrease in fucosylated structures (as detected by UEA-I), as malignancy increases—progressing from normal to adjacent and ultimately to tumor samples (**Figure 5B** and **Supplementary Figure 7B**).

**Figure 5.**
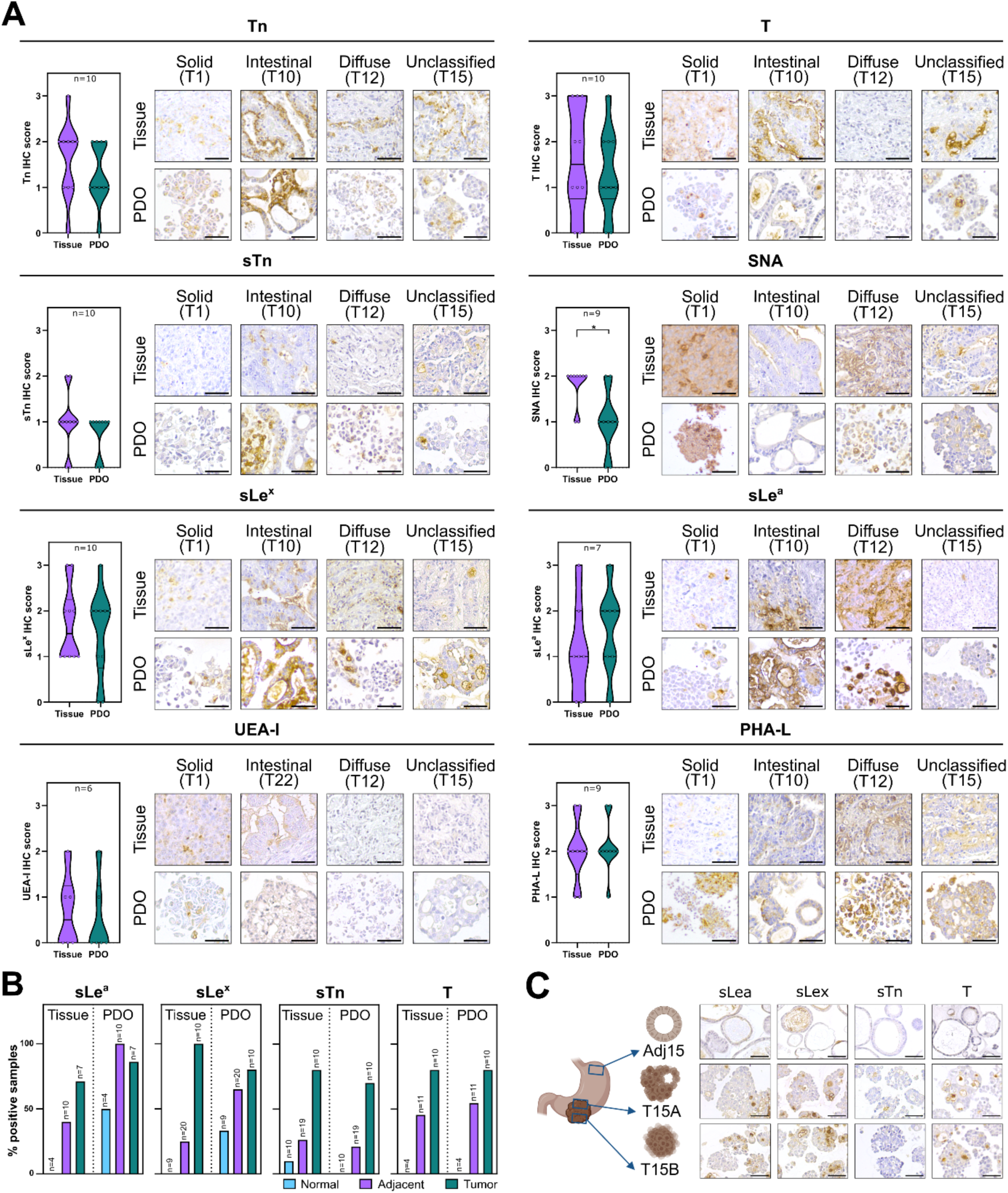
Gastric tumor PDOs retain the glycophenotype of parental tissues. **A)** Glycoprofiling of tumor tissues and corresponding tumor PDOs (T-PDOs) was evaluated, including short truncated *O*-glycans (T, Tn, and sTn) structures, α2,6-sialylated structures (SNA), sialylated type I Lewis (sLe^a^) and type II Lewis (sLe^x^) antigens, fucosylated structures (UEA-I), and branched *N*-glycans (PHA-L). Graphical quantification of glycan immunostaining of tumor tissues *versus* tumor PDOs (T-PDOs) is presented. Data represents the number of PDOs and tissues evaluated for each glycan. Representative images of the glycan profile of tissues and respective T-PDOs from different gastric cancer subtypes (intestinal, diffuse, unclassified mixed, and unclassified with solid pattern) are presented. *p value*<0.05 was considered statistically significant (unpaired t-test; * indicates a *p value*<0.05). **B)** Percentage of normal, adjacent and tumor samples, including both tissues and receptive PDOs, showing positive staining for sLe^a^, sLe^x^, sTn and T glycan antigens. **C)** Immunostaining for the sLe^a^, sLe^x^, sTn and T glycan antigens of samples derived from the same patient (Adj15 and the two tumor siblings PDOs, T15A and T15B). Scale bar corresponds to 50 µm.

To better understand the glycophenotype of both adjacent and tumoral epithelium components from the same patient, we evaluated the glycosylation profile on paired adjacent and sibling tumor PDOs derived from patients diagnosed with GC mixed unclassified subtype (patient T15 and T20) (**Supplementary Table 2**). Sibling T-PDOs from patient T15 originated two distinct histology: T-PDO 15A with a mixed glandular and solid pattern, and T-PDO 15B with a predominant solid pattern. On the other hand, sibling T-PDOs from patient T20 originated two T-PDOs with a well-defined histology with a non-cohesive phenotype (T-PDO 20A and 20B). T-PDOs clearly show a distinct glycoprofile when compared with Adj-PDOs, with increased staining for cancer-associated glycans such as sialylated Lewis antigens (sLe^a^ and sLe^x^), truncated *O*-glycans (sTn and T), Le^a^ or branched *N*-glycans (**Figure 5C** and **Supplementary Figure 7C-D**). Moreover, it was possible to observe that the predominance of some glycans in one sibling T-PDO compared with the other sibling T-PDO. For instance, T-PDO 15A is more positive for sTn and α2,6 sialylated *N*-glycans compared with T-PDO 15B, whereas the T-PDO 15B is more enriched in Lewis a and α2,3 sialylated structures (**Figure 5C** and **Supplementary Figure 7C**). This tumor heterogeneity was also found in the sibling T-PDOs from patient T20, where T20A is enriched on Lewis a and α2,6 sialylated *N*-glycans compared with T-PDO 20B, being negative for α2,6 sialylated structures (**Supplementary Figure 7D**). This analysis demonstrates the feasibility of using organoids as models to study the glycan epithelial heterogeneity between normal/adjacent and tumor-paired samples. Additionally, it highlights specific glycoprofile differences between cancer patients and within the same tumor specimen.

### Tumor PDO glycophenotype is preserved over time, biobanking and upon in vivo transplantation

The use of T-PDOs as a model to study glycan dynamics is dependent on their ability to keep their glycomic traits over time and passaging for *in vitro* and *in vivo* assays for mechanistic and functional studies, including drug testing. For that, T-PDO glycan heterogeneity was evaluated following continuous one-year culture, biobanking and freezing procedures, and xenotransplantation in mice models. We found that T-PDOs maintain their glycan heterogeneity in culture over time. Although a decrease in sTn and sLe^x^ expression was observed after 3 months in culture (approximately 12 passages), the positivity of these glycans was maintained for over 12 months in culture (approximately 48 passages) (**Figure 6A** and **Supplementary Figure 8A**). Importantly, when subjected to biobanking and subsequent re-culturing, no major differences were observed in glycan expression patterns (**Figure 6B** and **Supplementary Figure 8B**), indicating that this is a better approach to preserve T-PDO glycan phenotypic features. Then, five PDOX were successfully established from T-PDOs with different types of histology (solid: T1, T1.1; glandular: T10, T22; unclassified mixed: T15B) (**Figure 1D**, **Supplementary Figure 2B**; and **Supplementary Table 2**). T-PDOs with non-cohesive phenotype were not able to form a tumor in mice (**Supplementary Table 2**). We observed that tumor tissues derived from PDOX maintained the glycan expression phenotype and heterogeneity (**Figure 6C**) mimicking the corresponding T-PDO and original human tumor tissue glycan profile and architecture (**Figure 5**). These data demonstrate that T-PDOs can be used as a model to study and test glycan-related drugs in *in vivo* pre-clinical settings. Then, to evaluate if organoids derived from patient xenotransplanted mice (PDXs) also retain the glycophenotype of the original tumor tissues, two PDX-Organoids (PDXO T19 and T30) were generated (**Figure 6D**, **Supplementary Figure 2B**, and **Supplementary Table 2**). Although both PDX and PDXOs retain the original histologic intestinal GC subtype (**Supplementary Figure 2B**), the PDXOs did not reflect the glycan profile of the original human tumor, being almost depleted for glycans with the exception of the truncated *O*-glycan T, and *de novo* expression of α2,6-sialylated *N-*glycans (SNA) and branched *N*-glycans (PHA-L) (**Figure 6D**). We postulate that these observations are most likely due to a clonal selection of tumor cells when grown in the mice or due to different glycan machinery present in mice compared with humans.

**Figure 6.**
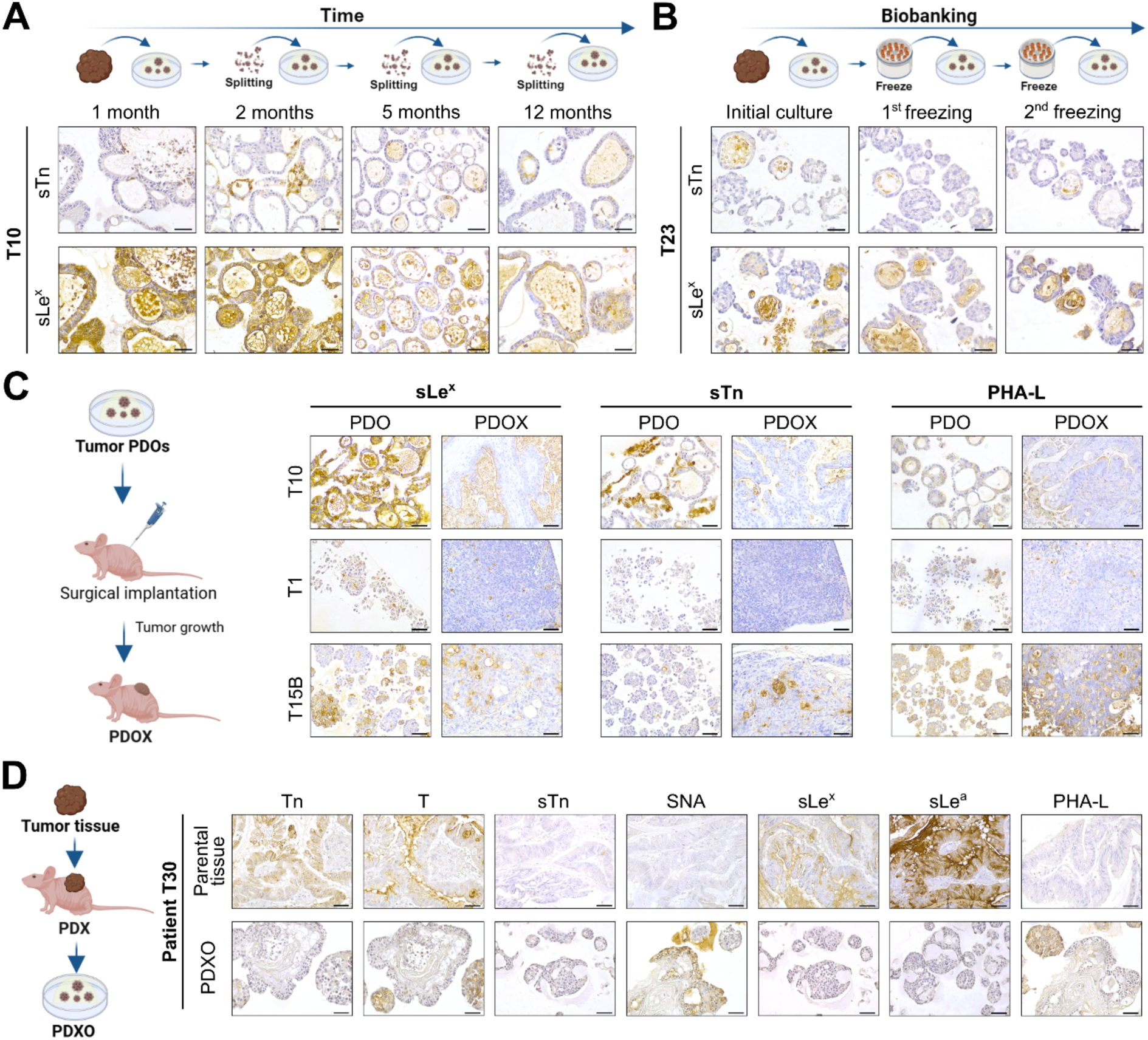
Tumor PDOs can maintain their glycan heterogeneity in culture over time, upon biobanking and *in vivo* xenografting. **A)** sTn and sLe^x^ expression analysis up to 1-year continuous culture. **B)** sTn and sLex expression analysis upon biobanking and re-culture of T-PDOs. **C)** PDOXs mimic the glycan landscape of corresponding PDOs. **D)** Patient-derived xenograft organoids (PDXOs) mimic the histologic features but not the glycoprofile of the parental human tissues. Scale bar corresponds to 50 µm.

## DISCUSSION

Aberrant glycosylation plays a crucial role in gastric carcinogenesis and is a key feature actively dictating multiple tumor aggressive features^2, 3^. However, due to the complexity and dynamics of this cellular process, there is a need to use advanced disease models to study glycans throughout the gastric carcinogenic cascade. In this work, we have shown the significance of PDOs as a valuable *ex vivo* model to study complex glycosylation patterns within normal, pre-malignant, and tumoral human gastric epithelium. This study bridges an existing knowledge gap concerning the glycosylation patterns within these 3D models and their implications in gastric pathophysiology.

We generated a gastric PDO biobank comprising 56 PDOs derived from non-malignant, pre-malignant, and tumoral tissues including early and later cancer stages of different GC subtypes which can be subject to further studies of GC biology and tumor progression. Other GC PDO biobanks have been reported, however, only one described PDOs derived from cancer-free individuals^32, 33, 44–46^. Indeed, one of the key observations of this study relates to the morphological heterogeneity exhibited by gastric PDOs derived from cancer patient’s adjacent mucosa. Normal gastric PDO morphology is described as cystic, spherical, and composed of a single layer of cells with a central lumen^27, 36^. The different morphologies observed in this study raise the question of whether these organoids might recapitulate various stages of normal gastric epithelial differentiation or gastric carcinogenesis. While all normal PDOs consistently expressed gastric mucins (MUC5AC and MUC6) along with other markers associated with distinct gastric cellular compartments, the identification of organoids expressing markers associated with pre-malignant conditions stands out as a particularly relevant finding. The morphological characteristics and the retention of IM markers (MUC2 and CDX2), and the concomitant increased levels of Ki-67 and CEACAM5 suggest that PDOs recapitulate the early stages of carcinogenesis, including IM and dysplasia. This is in agreement with previous studies that have described that non-tumoral gastric PDOs may express the IM marker CDX2^46^. Moreover, it was also described that CDX2 overexpression in gastric organoids can induce incomplete IM^47^. More recently, was shown that around 80% of PDOs derived from gastric tissues with IM condition express several IM molecular markers similar to the ones present in gastric IM and cancer tissues^48^. Additionally, PDOs derived from stem cells isolated from gastric dysplastic lesions showed morphologic alterations characterized by multilayering of cells and disorganized nuclei^49^. Together, these findings suggest that careful characterization should be considered when using PDOs derived from tissues adjacent to the tumor when studying normal physiologic processes of the gastric epithelium. As such, our established biobank of PDOs derived from cancer-free patients is a valuable addition to the scientific research community to study not only non-cancer-related pathologies, but importantly pre-malignant gastric cancer conditions.

In this work, we validated the glycoprofile of normal and pre-malignant PDOs as avatars of the *in vivo* original tissues. We found that PDOs mimic the glycoprofile associated with the normal gastric mucosa, as well as the tissue mucosa displaying inflammation-related markers (such as sLe^x^ and sLe^a^)^12, 13^ and IM/dysplasia (including the truncated O-glycan sTn)^42^ resembling the findings from original tissues characterized by chronic gastritis and IM lesions. Moreover, the decrease in UEA-I staining alongside an increase in sLe^x^ expression in Adj-PDOs suggests that H2 antigen (detected by UEA-I lectin) may be consumed through sialylation and α1,3-fucosylation, promoting sLe^x^ biosynthesis. This shift is likely driven by altered glycosyltransferase activity and may represent a potentiating mechanism for malignancy-associated glycan remodeling. Indeed, we observed that sialylated type I and II Lewis antigens are expressed preferentially in undifferentiated PDOs, suggesting that both sLe^a^ and sLe^x^ may have a role in the maintenance of stem-like cellular phenotypes^50^. Our study also demonstrated that type I and II Lewis antigens can be modulated according to PDOs differentiation status and impact the binding capabilities of *H. pylori* to gastric cells, a major contributor to gastric inflammation and carcinogenesis^5, 6^. We show that 17875/Leb *H. pylori* strain has higher binding capacity in the differentiated foveolar pit-type PDOs when compared with mixed-gland PDOs, which reflects an increased expression of the Le^b^ molecular anchor. In agreement, Aguilar, *et al*^31^ also observed a higher binding of *H. pylori* in more differentiated foveolar PDOs^31, 51^. Although the PDO glycosylation profile was not characterized in their study, the *H. pylori* strain used in their work also binds to Le^b^-glycoconjugates, P12^52^, which is in line with our findings. Here, we also demonstrate that the SabA-positive *H. pylori* strain binds preferentially to PDOs derived from adjacent tissues and well as to mixed-gland PDOs, that express sLe^x/a^, when compared with PDOs from normal gastric tissues or with differentiated foveolar PDOs. These data indicate that the complexity of PDOs glycosylation profile can offer crucial insights into the glycan-dependent interactions underpinning *H. pylori* colonization and pathogenicity.

From a clinical perspective, the limited availability of fresh clinical samples makes the analysis of specific glycan landscapes in the epithelial component extremely difficult, especially due to the contamination by stromal and immune cells. This work showed the significant fidelity of T-PDOs in replicating the histology and glycan profiles of tumor tissues from distinct GC histologic subtypes. Our data is in agreement with a previous study showing that organoids derived from pancreatic cancer PDXs retain the *N-*glycosylation diversity and relative abundance observed in the matched PDX model^53^. Moreover, the T-PDO glycophenotypes are preserved when subjected to biobanking, extended culture periods, and after xenografting procedures. Despite the observed loss of sTn and sLe^x^ expression in tumor PDOs upon 5 months of continuous culture, our findings support previous studies demonstrating genomic and transcriptomic shifts in PDOs after long-term culture (3 months for non-tumoral PDOs and 6 months for T-PDOs)^27, 32^. This loss of heterogeneity may be a result of the selective processes over passage, and raise the the possibility of certain subpopulations of tumor cells with distinct glycomic profiles and/or best survival capabilities being favored during long-term culture conditions.

Here, we demonstrate the utility of using PDOs as a model to study both non-tumoral and tumoral epithelial glycome. By using tumor PDOs derived from different tumor regions and paired normal organoids, we were not only able to elucidate the intra- and inter-glycan heterogeneity, having a more comprehensive overview of the tumor landscape, but also to identify specific glycan antigens restricted to/enriched in both adjacent and different tumoral regions. Our combined approach with “sibling” tumor PDOs and paired adjacent PDOs holds the potential to uncover previously elusive molecular markers with diagnostic and prognostic relevance, as well as to support the development of novel personalized therapies. Indeed, in a recent study, we demonstrated the use of T-PDOs as a pre-clinical tool to validate monensin as a potential repurposing drug against sLe^x^-positive gastric tumors^37^.

In summary, this groundbreaking study explored the complex glycosylation phenotypes in normal, adjacent, and tumor PDOs, and highlights the potential of harnessing the power of PDOs as a model to study gastric carcinogenesis. Our results pave the way for further research into the functional implications of aberrant glycosylation in gastric carcinogenesis, positioning PDOs as a valuable tool to dissect the molecular mechanisms underlying cancer initiation and progression. Furthermore, the interactions between glycans, non-epithelial cells, and extracellular matrix components represent an important area for future investigation. Overall, this study establishes PDOs as a robust *ex vivo* model to explore glycosylation within the broader context of gastric pathophysiology.

## Acknowledgments

We thank all the patients for participating in this study. The authors thank Dina Leitão from the Service of Anatomy Pathology of Unidade Local de Saúde (ULS) de São João, Porto, Portugal, for the support in the optimization of the CDX2 immunohistochemistry. The authors acknowledge the support of the i3S Scientific Platforms: Advanced Light Microscopy, Animal Facility, BioSciences Screening, Cell Culture and Genotyping, and Histology and Electron Microscopy, members of the PPBI (PPBI-POCI-01-0145-FEDER-022122). We thank the support of Horizon 2020 WIDESPREAD-05–2018-TWINNING Remodel under the EU grant agreement no. 857491.

## Data Availability

The data, analytical methods and study materials are available to other researchers by request.

