## Supplementary for "Patient-derived organoids to study glycosylation dynamics during gastric disease"

Liliana Santos-Ferreira *et al.*

**Supplementary Table 1 -** Clinicopathological data of obese and gastric cancer patients used for the development of non-tumoral patient-derived organoids.

| Disease status | Patient/Tissue code | PDO | Gender | Age | Patient clinical diagnose | PDO-derived mucosa histology | PDO Histology |
|---|---|---|---|---|---|---|---|
| Obese - Normal gastric mucosa | N1 | N1 | M | 52 | No significant alterations | Normal | Normal |
| | N2 | N2 | F | 60 | ChrGastritis | Gastritis | Normal |
| | N3 | N3 | M | 56 | ChrGastritis; IM | Gastritis/IM | Normal/Complex (16-20%) |
| | N4 | N4 | F | 49 | ChrGastritis | Normal | Normal |
| | N5 | N5 | F | 49 | ChrGastritis | Gastritis | Normal |
| | N6 | N6 | F | 42 | ChrGastritis | Normal | Normal/Complex (<5%) |
| | N7 | N7 | F | 49 | ChrGastritis; IM | Normal | Normal |
| | N8 | N8 | F | 56 | ChrGastritis; IM | Normal | Normal |
| | N9 | N9 | M | 46 | ChrGastritis | Gastritis | N.A. |
| | N10 | N10 | M | 62 | ChrGastritis | Gastritis | Normal |
| | N11 | N11 | F | 48 | ChrGastritis | Normal | Normal |
| Gastric cancer - Tumor adjacent gastric mucosa | Adj1 | Adj1 | F | 71 | ChrGastritis; IM | N.A. | Normal/Complex (<5%) |
| | Adj3 | Adj3 | M | 71 | ChrGastritis; IM | Gastritis; IM | Normal/Complex (5-10%) |
| | Adj4 | Adj4 | M | 68 | ChrGastritis; IM | Gastritis | Normal |
| | Adj5 | Adj5 | F | 86 | ChrGastritis; IM | Gastritis | Normal |
| | Adj6 | Adj6 | M | 77 | ChrGastritis; IM | Gastritis; IM | Normal |
| | Adj7 | Adj7 | M | 70 | ChrGastritis; IM | Gastritis | N.A. |
| | Adj9 | Adj9 | M | 68 | ChrGastritis; IM | Gastritis; IM | Normal/Complex (5-10%) |
| | Adj11 | Adj11A<br>Adj11B | F | 55 | No significant alterations | Gastritis | Adj11A – Normal<br>Adj11B - Normal/Complex (11-15%) |
| | Adj13 | Adj13 | M | 68 | IM; Dysplasia | Gastritis; IM | Normal/Complex (5-10%) |
| | Adj14 | Adj14 | M | 60 | ChrGastritis; IM | Gastritis | Normal |
| | Adj15 | Adj15 | F | 78 | ChrGastritis; IM | Gastritis | Normal/Complex (<5%) |
| | Adj16 | Adj16A<br>Adj16B | M | 62 | ChrGastritis; IM | Gastritis | Adj16A – Normal<br>Adj16B - Normal/Complex (<5%) |
| | Adj17 | Adj17 | F | 67 | ChrGastritis; IM | Gastritis; IM | Normal/Complex (11-15%) |
| | Adj18 | Adj18 | F | 77 | ChrGastritis; IM | Gastritis; IM | Normal/Complex (16-20%) |
| | Adj19 | Adj19 | M | 61 | ChrGastritis; IM | Gastritis; IM | Normal/Complex (16-20%) |
| | Adj20 | Adj20 | M | 51 | ChrGastritis; IM | Gastritis | Normal/Complex (5-10%) |
| | Adj21 | Adj21 | F | 54 | ChrGastritis; IM | Gastritis; IM | Normal |
| | Adj22 | Adj22A<br>Adj22B | F | 80 | ChrGastritis; IM | Gastritis; IM | Adj22A – Normal<br>Adj22B – Normal/Complex (5-10%) |
| | Adj23 | Adj23 | F | 58 | ChrGastritis; IM | Gastritis; IM | Normal |
| | Adj25 | Adj25 | F | 76 | ChrGastritis; IM | Gastritis | Normal |
| | Adj26 | Adj26 | M | 78 | ChrGastritis; IM | Gastritis | Normal/Complex (5-10%) |
| | Adj27 | Adj27 | M | 62 | ChrGastritis; IM | Normal | Normal/Complex (<5%) |
| | Adj29 | Adj29 | M | 88 | ChrGastritis; IM | Gastritis; IM | Normal/Complex (5-10%) |

**ChrGastritis**– Chronic Gastritis; **IM** – Intestinal Metaplasia**; N.A.** – Non available material; **Normal structures:** single cell layer with a central lumen; ***Complex structures:** complex layer of cells with/without cribriform or vacuole-like structures; **Grey highlight** – sibling PDOs.

**Supplementary Table 2 -** Clinicopathological data of gastric cancer patients used to develop tumoral patient-derived organoids and receptive histology regarding PDOs culture and PDOX tissue.

| Patient/Tissue code | PDO | Gender | Age | Lauren | WHO | T | N | M | Stage | PDO derived tumor tissue Histology | PDO Histology | Xenograft Histology |
|---|---|---|---|---|---|---|---|---|---|---|---|---|
| **T1** | T1 | M | 71 | Unclassified | Tubular (Poorly differentiated) | T4a | N1 | M0 | IIIA | Solid pattern | Solid | Solid |
| | T1.1 | | | | | | | | | | Solid | Solid |
| **T6** | T6 | M | 77 | Unclassified | Mixed | T4a | N2 | M0 | IIIB | Intestinal | Solid + Glandular | N.A. |
| **T10** | T10 | F | 89 | Intestinal | Tubular | T1a | N0 | M0 | IA | Intestinal | Glandular* | Intestinal |
| **T12** | T12A | M | 64 | Diffuse | Poorly cohesive | T4a | N2 | M0 | IIIB | Diffuse (Poorly cohesive w/ a glandular component) | Non-cohesive cells | N.A. |
| | T12B | | | | | | | | | | Non-cohesive cells | No tumor observed |
| **T15** | T15A | F | 78 | Unclassified | Mixed (Tubular+ Solid) | T3 | N0 | M0 | IIA | Intestinal with solid pattern | Glandular + Solid | N.A. |
| | T15B | | | | | | | | | | Glandular + Solid | Intestinal with solid pattern |
| **T19** | T19 | M | 61 | Intestinal | Tubular | T3 | N1 | M0 | IIB | Intestinal with solid pattern | Glandular + Solid | Intestinal# |
| **T20** | T20A | M | 51 | Unclassified | Mixed (>90% non-choesive cells) | T3 | N3a | M0 | IIIB | Diffuse (Poorly cohesive w/ mucinous component) | Non-cohesive cells + Solid | No tumor observed |
| | T20B | | | | | | | | | | Non-cohesive cells + Solid | No tumor observed |
| **T21** | T21A | F | 54 | Diffuse | Poorly cohesive | T3 | N2 | M0 | IIIA | Diffuse (Poorly cohesive w/ a glandular component) | Glandular | N.A. |
| | T21B | | | | | | | | | | Non-cohesive cells + Solid | N.A. |
| **T22** | T22A | F | 80 | Unclassified | Mixed (Tubular + poorly cohesive cells) | T4a | N1 | M0 | IIIA | Intestinal | Glandular | No tumor observed |
| | T22B | | | | | | | | | | Glandular | Intestinal |
| **T23** | T23 | F | 58 | Intestinal | Tubular | T1b (m) | N0 | M0 | IA | Intestinal | Glandular | No tumor observed |
| **T27** | T27 | M | 62 | Unclassified | Mixed | T4a | N0 | M1 | IV | Intestinal w/ poorly cohesive component | Non-cohesive cells | N.A. |
| **T29** | T29 | M | 88 | Unclassified | Mixed (Tubular w/mucinous component) | T2 | N0 | M0 | IB | Intestinal w/ poorly cohesive component | Glandular | No tumor observed |
| **T30** | T30 | M | 75 | Intestinal | Tubular | T1 | N0 | M0 | IA | Intestinal | Glandular + Solid | Intestinal with solid pattern# |

*with cribriform structures

#PDOs derived from patient-derived xenografts

**Grey highlight** – sibling PDOs.

**Supplementary Table 3** - Primers used in this work for qPCR analysis.

| Gene | Forward primer | Reverse primer | PCR product size (bp) |
|---|---|---|---|
| *MUC5AC* | CTTCTCAACGTTTGACGGGAAGC | CTTGATCACCACCACCGTCTG | 190 |
| *MUC6* | GCCCCGGTATCTTCTCTCGG | ACACCTGCAGGGTGAGTACG | 283 |
| *MUC2* | GACCTCCAGCACAGTTTTATCAACA | GCCAGCAACAATTGACACGTATCT | 305 |
| *CDX2* | AACCAGGACGAAAGACAAAT | GAAGACACCGGACTCAAG | 311 |
| *ACTB* | TGCTATCCAGGCTGTGCTAT | AGTCCATCACGATGCCAGT | 62 |
| *PGC* | AGAGCCAGGCCTGCACCAGT | GCCCCTGTGGCCTGCAGAAG | 580 |
| *STT* | CTAGAGTTTGACCAGCCAC | GACAGATCTTCAGGTTCCAG | 268 |

**Supplementary Table 4 -** List of antibodies and lectins used in this study.

| Antigens | Clone/Lectin | Method (dilution) | Host and Isotype | Retrieval | Supplier/ Reference |
|---|---|---|---|---|---|
| MUC5AC | 45M1 | IHC (1:1000)<br>IF (1:1000) | Mouse IgG | 10 min | [1] |
| MUC6 | CLH5 | IHC (1:100)<br>IF (1:100) | Mouse IgG1 | 10 min | Santa Cruz Biotechnology |
| PGC | D-5 | IHC (1:150) | Mouse IgG2a | 20 min | Santa Cruz Biotechnology |
| H/K+ ATPase | C-4 | IHC (1:2000) | Mouse IgG2b | 20 min | Santa Cruz Biotechnology |
| Ki-67 | D2H10 | IHC (1:500) | Rabbit IgG | 40 min | Cell Signalling |
| CEACAM5 | CB30 | IHC (1:500) | Mouse IgG1 | No Retrieval | Cell Signalling |
| CDX2 | EPR2764Y | IHC (undiluted; Ventana BenchMark ULTRAStaining System) | Rabbit | 40 min | Roche/Ventana Medical Systems |
| sTn | B72.3 | IHC (1:5) | Mouse IgG1 | No Retrieval | [2] |
| sLe$^{x}$ | CSLEX1 | IHC (1:250)<br>IF (1:200) | Mouse IgM | No Retrieval | BD Pharmingen |
| Le$^{a}$ | Ca3F4 | IHC (1:5) | Mouse IgG2a | 10 min | [3] |
| Le$^{b}$ | BG-6 (T218) | IHC (1:50)<br>IF (1:50) | Mouse IgM | 10 min | Signet |
| sLe$^{a}$ | CA19.9 | IHC (1:500)<br>IF (1:500) | Mouse IgG | 15 min (IF)<br>No Retrieval (IHC) | Abcam |
| Le$^{y}$ | AH6 | IHC (1:10) | Mouse IgM | 10 min | [4] |
| Le$^{x}$ | SH1 | IHC (1:5)<br>IF (1:5) | Mouse IgG3 | 15 min | [5] |
| Tn | 5F4 | IHC (1:10) | Mouse IgM | 20 min | [7] |
| T | 3C9 | IHC (1:10) | Mouse IgM | 20 min | [8] |
| a. Type 2 blood group H; b. Fucα1-2Galb1-4Glc; c. Le$^{y}$ | UEA-I | Lectin-HC (1:1000) | - | No Retrieval | Vector Labs |
| a. α2-6 sialylated LacNAc; b. α2-6-sialylated LacdiNAc | SNA | Lectin-HC (1:750) | - | No Retrieval | Vector Labs |
| B1-6 branched *N*-glycans | PHA-L | Lectin-HC (1:400) | - | No Retrieval | Vector Labs |
| Bisecting GlcNAc with Type 2 LAcNAc | PHA-E | Lectin-HC (1:600) | - | No Retrieval | Vector Labs |
| a. Terminal 3-*O*-sulfated Gal on LacNAc*;* b. α2-3-sialylated LacNAc | MAL-I | Lectin-HC (1:1000) | - | No Retrieval | Vector Labs |
| **Secondary antibodies** | **Conjugates** | **Method (dilution)** | **Host and Isotype** | **Retrieval** | **Supplier/ Reference** |
| Anti-Rabbit Immunoglobulins | Biotinylated | IHC (1:200) | Swine Polyclonal | - | Dako Denmark (E0353) |
| Anti-Mouse Immunoglobulins | Biotinylated | IHC (1:200) | Rabbit Polyclonal | - | Dako Denmark (E0354) |
| Anti-mouse | Alexa Fluor 488 | IF (1:500) | Goat IgG | - | Invitrogen (A11029)) |
| Anti-mouse | Alexa Fluor 594 | IF (1:500) | Goat IgM | - | Invitrogen (A21044) |

**IHC**– Immunohistochemistry; **IF** – Immunofluorescence; **Lectin-HC** – Lectin-based histochemistry.

**SUPPLEMENTARY DATA_SUPPLEMENTARY FIGURES**

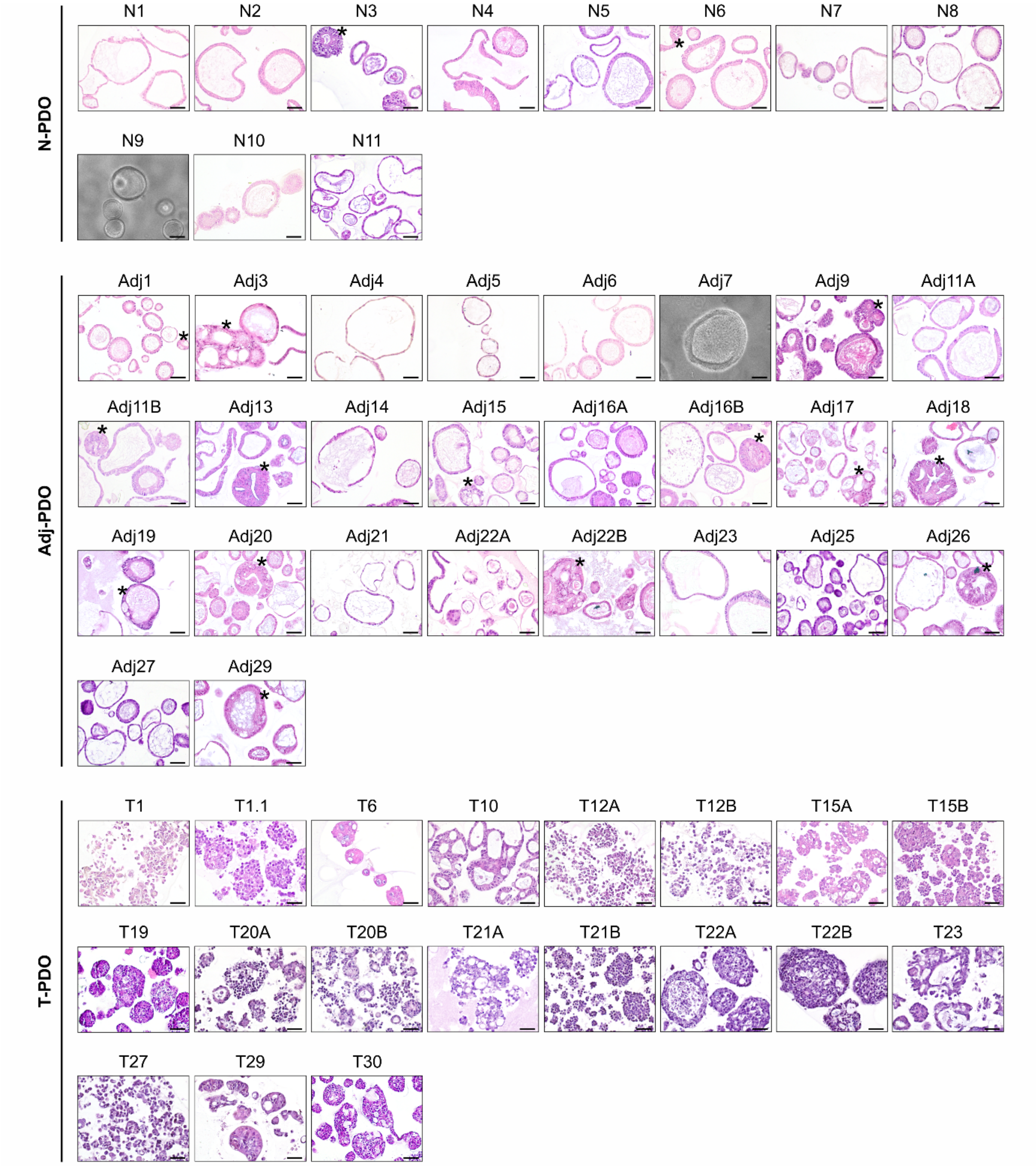


**Supplementary Figure 1. Gastric patient-derived organoid biobank overview.** Representative hematoxylin and eosin staining (H&E) and brightfield (BF) images of PDOs derived from non-cancer gastric mucosa (N), gastric mucosa adjacent to the tumor (Adj) and gastric tumor (T) tissues available in our biobank. *, indicates PDOs with complex morphologies mimicking premalignant-like conditions. Scale bars correspond to 50 µm.

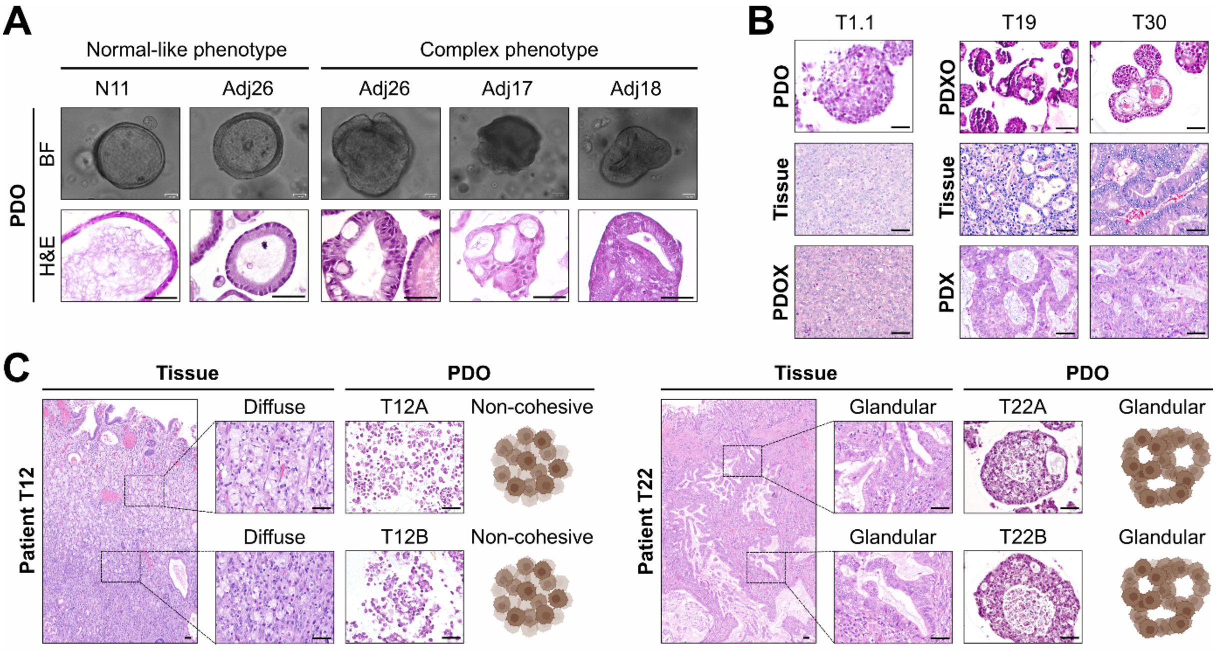


**Supplementary Figure 2. Morphologic and histologic features of gastric patient-derived organoids. A)** Representative BF and H&E images of PDOs derived from non-tumoral PDOs (N-PDO 11, and Adj-PDO 17, 18, and 26 are represented) with normal-like and complex phenotypes**. B)** Representative H&E images of tumor PDO (T-PDO), respective parental tissue, and respective patient-derived organoid xenograft (PDOX). Two tumor patient-derived xenograft organoids (PDXO) and their respective human and mice xenograft tissue (PDX) are represented. T-PDOs replicate the histologic features of respective tissues. **C)** "Sibling" T-PDOs derived from different tumor fragments of patient T12 (right panel) and patient T22 (left panel) diagnosed with diffuse and intestinal subtypes, respectively. Scale bars correspond to 50 μm.

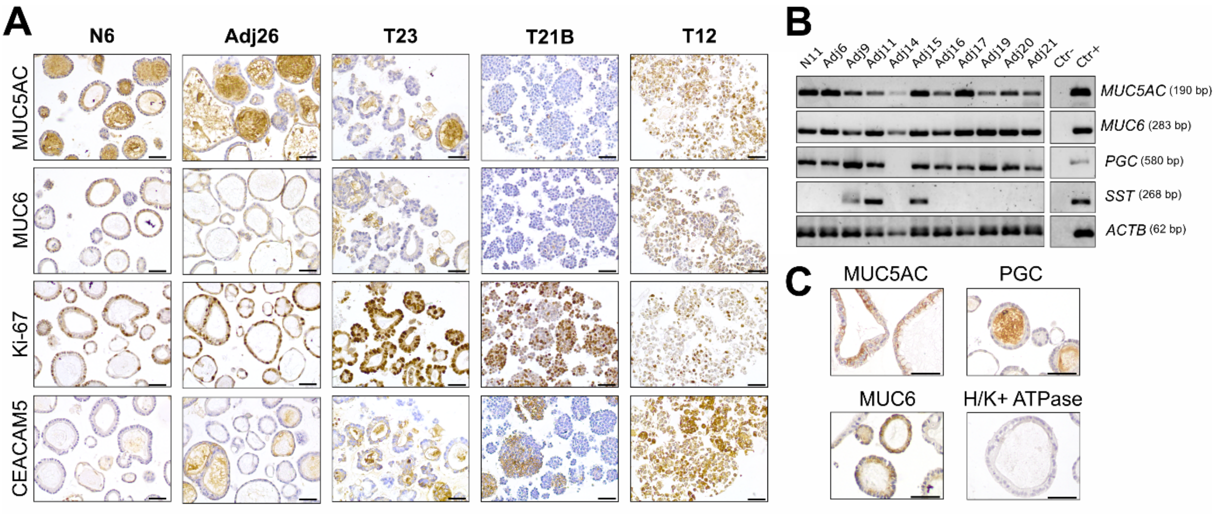


**Supplementary Figure 3. Normal and premalignant gastric patient-derived organoids characterization. A)** Representative immunohistochemistry images of normal, adjacent and tumor PDOs showing staining of the gastric classical mucins MUC5AC and MUC6, the proliferation marker Ki-67, and the tumoral biomarker CEACAM5. **B)** Characterization of normal and adjacent PDOs showing the expression of gastric epithelial differentiation markers, *MUC5AC*, *MUC6*, pepsinogen C (*PGC*), and somatostatin (*SST*). *ACTB* was used as the housekeeping control gene. **C)** Representative immunohistochemistry images of non-tumoral PDOs for the staining of gastric markers (MUC5AC, MUC6, PGC, and H/K+ ATPase). Scale bars correspond to 50 µm.

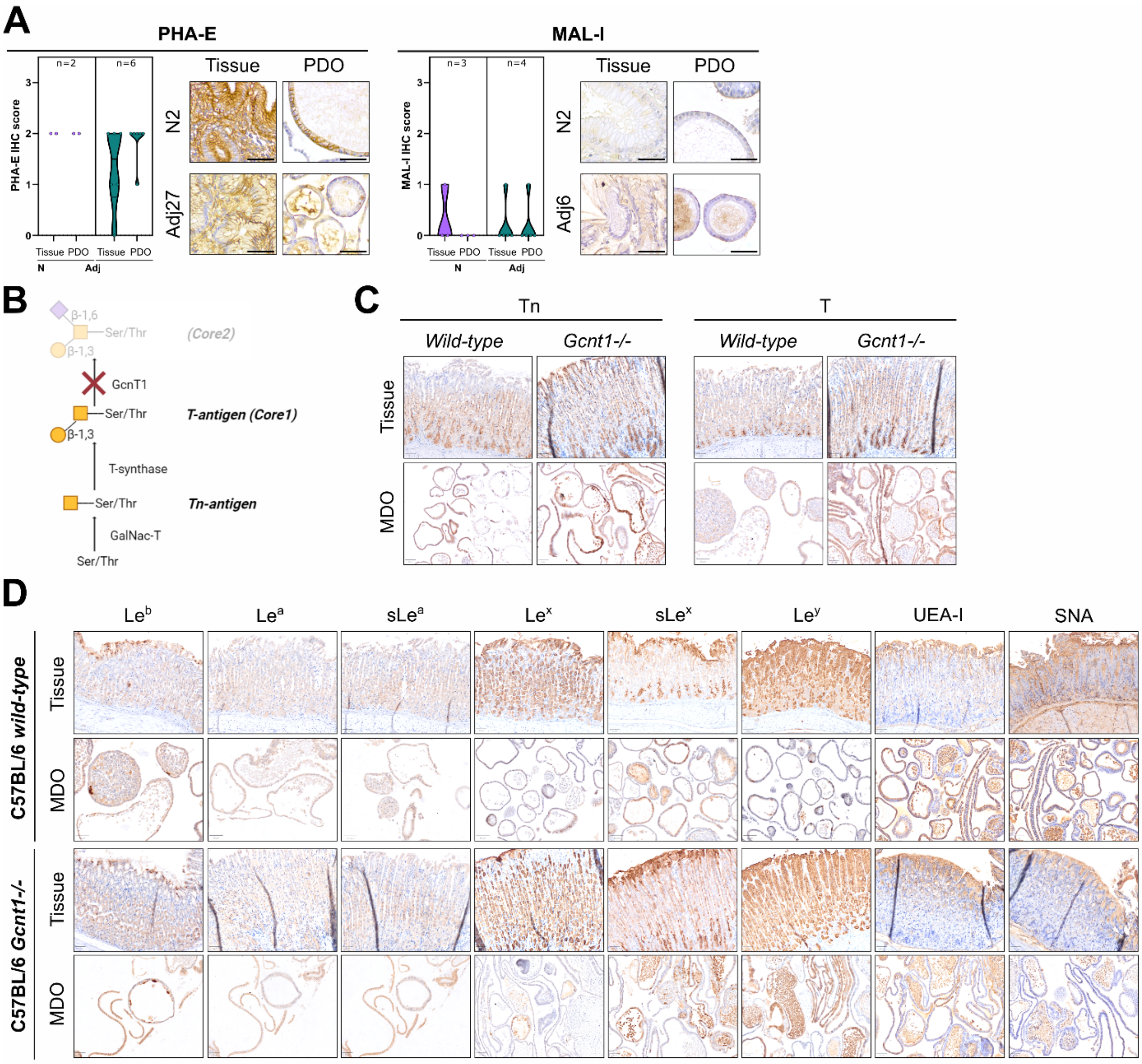


**Supplementary Figure 4. Non-tumoral gastric human and mouse PDOs mimic the glycophenotype of parental tissues. A)** Quantification of the imunnostaining for bisecting *N*-glycans (Phaseolus vulgaris erythroagglutinin - PHA-E), and α2,3 sialylated LacNAc (Maackia amurensis lectin I - MAL-I). Representative pictures of the glycoprofile found in normal (N) and adjacent (Adj) PDOs and matched tissues are also presented. **B)** Schematic representation of the glycoprofile synthesis of C57BL/6 *wild-type* and *Gcnt1-/-* mice. **C)** Immunostaining for the core 1 glycans (T and Tn) showing an enrichment on the KO (*Gcnt1-/-*) model both in the tissue and in the mouse-derived organoids (MDO) when compared with the *wild-type* model. **D)** Representation of the glycoprofile found in C57BL/6 *wild-type* and *Gcnt1-/-* MDOs and matched tissues. Type I Lewis antigens ($Le^b$, $Le^a$, $sLe^a$), type II Lewis antigens ($Le^y$, $Le^x$, $sLe^x$), α2,3 sialylated (Sambucus Nigra Lectin - SNA), and fucosylated (Ulex Europaeus Agglutinin I - UEA-I) glycans were analyzed. Scale bars correspond to 50 µm.

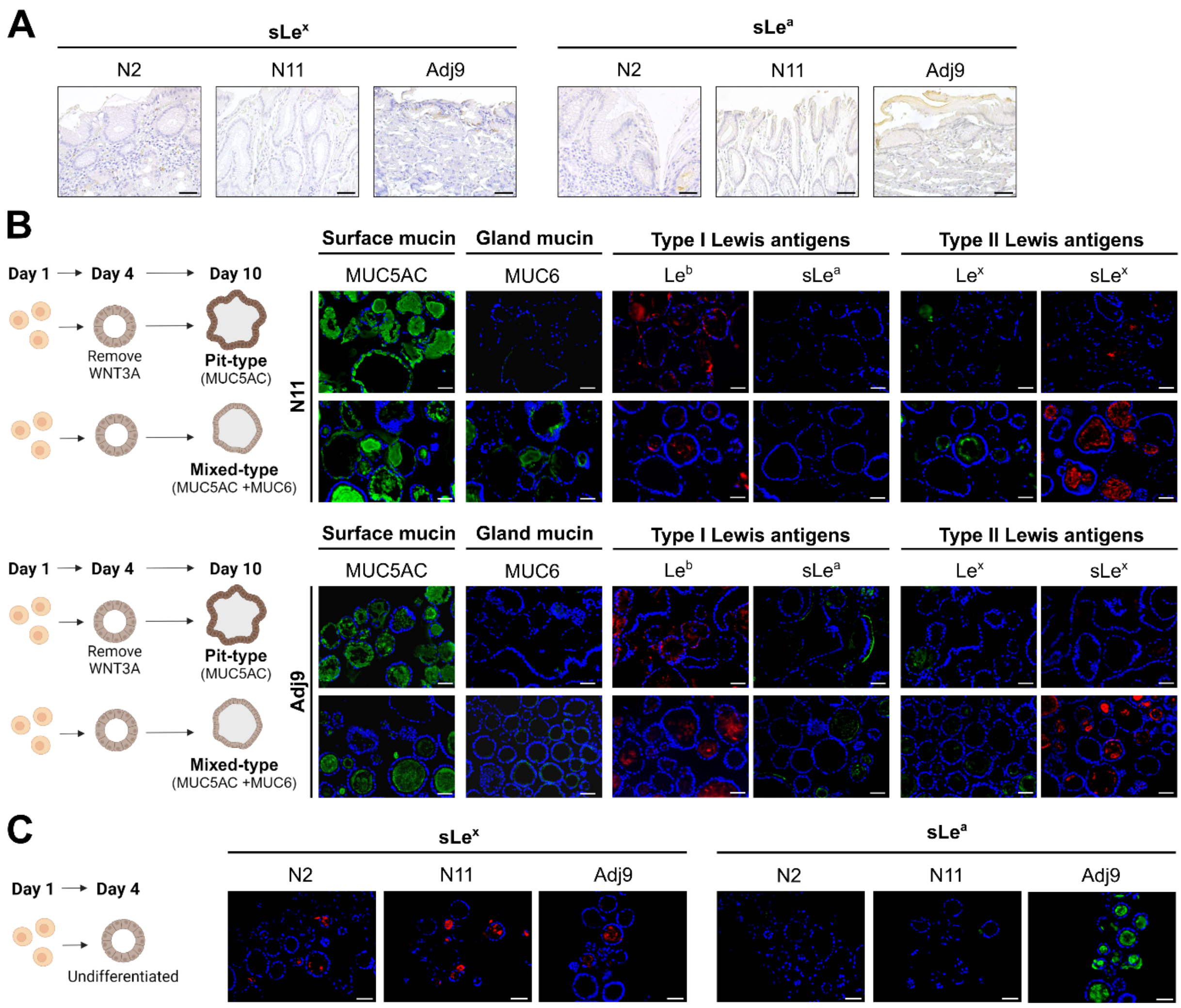


**Supplementary Figure 5. PDO type I/II Lewis glycophenotype depends on foveolar and mucous gland differentiation. A)** Immunostaining images for sialylated Lewis antigens (sLe$^x$ and sLe$^a$) in the parental tissues from where PDOs were originated, showing negative or focal expression of these antigens. **B)** PDOs differentiation on foveolar pit-type and gland mixed-type and their surface (MUC5AC) and gland (MUC6) mucin, and type I and II Lewis expression patterns on two additional PDOs (N11 and Adj9). **C)** Undifferentiated normal and adjacent PDOs express high levels of sialylated type I and type II Lewis antigens (sLe$^a$ and sLe$^x$).

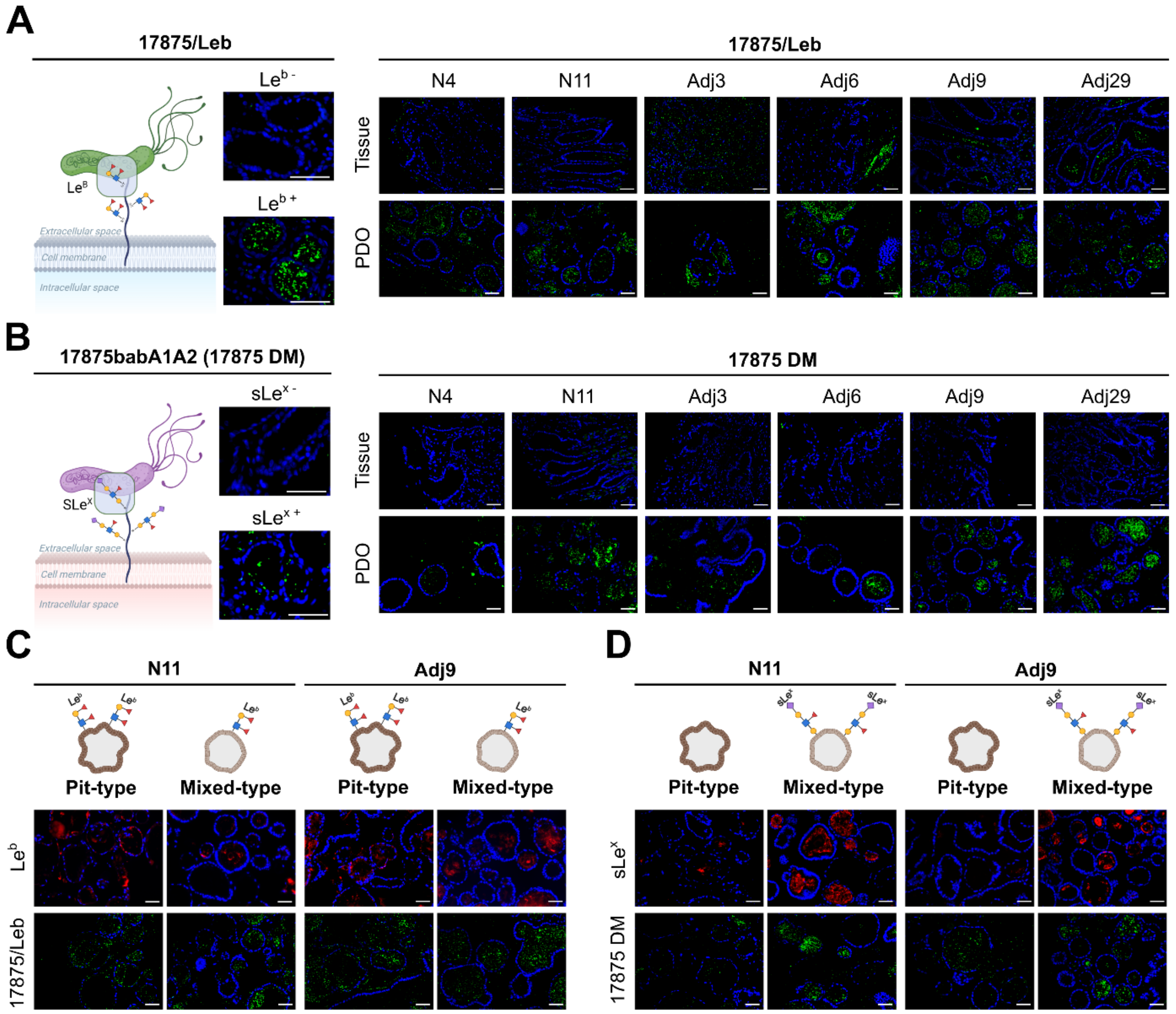


**Supplementary Figure 6. PDO type I/II Lewis glycophenotype influences *Helicobacter pylori* binding. A)** Schematic representation of the 17875/Leb *H. pylori* strain binding to the Le$^{b}$ antigen with representative negative and positive controls. Representative images of 17875/Leb *H. pylori* binding in N4, N11, Adj3, Adj6, Adj9, and Adj29 tissues and respective PDOs. **B)** Schematic representation of the 17875 DM *H. pylori* strain binding to the sLe$^{x}$ antigen with representative negative and positive controls. Representative images of 17875 DM *H. pylori* binding in N4, N11, Adj3, Adj6, Adj9, and Adj29 tissues and respective PDOs. **C)** Binding assay of 17875/Leb strain to N11 and Adj9 pit-type and mixed-type and corresponding Le$^{b}$ expression. **D)** Binding assay of 17875 DM strain to N11 and Adj9 pit-type and mixed-type and corresponding sLe$^{x}$ expression. Scale bar corresponds to 50 µm.

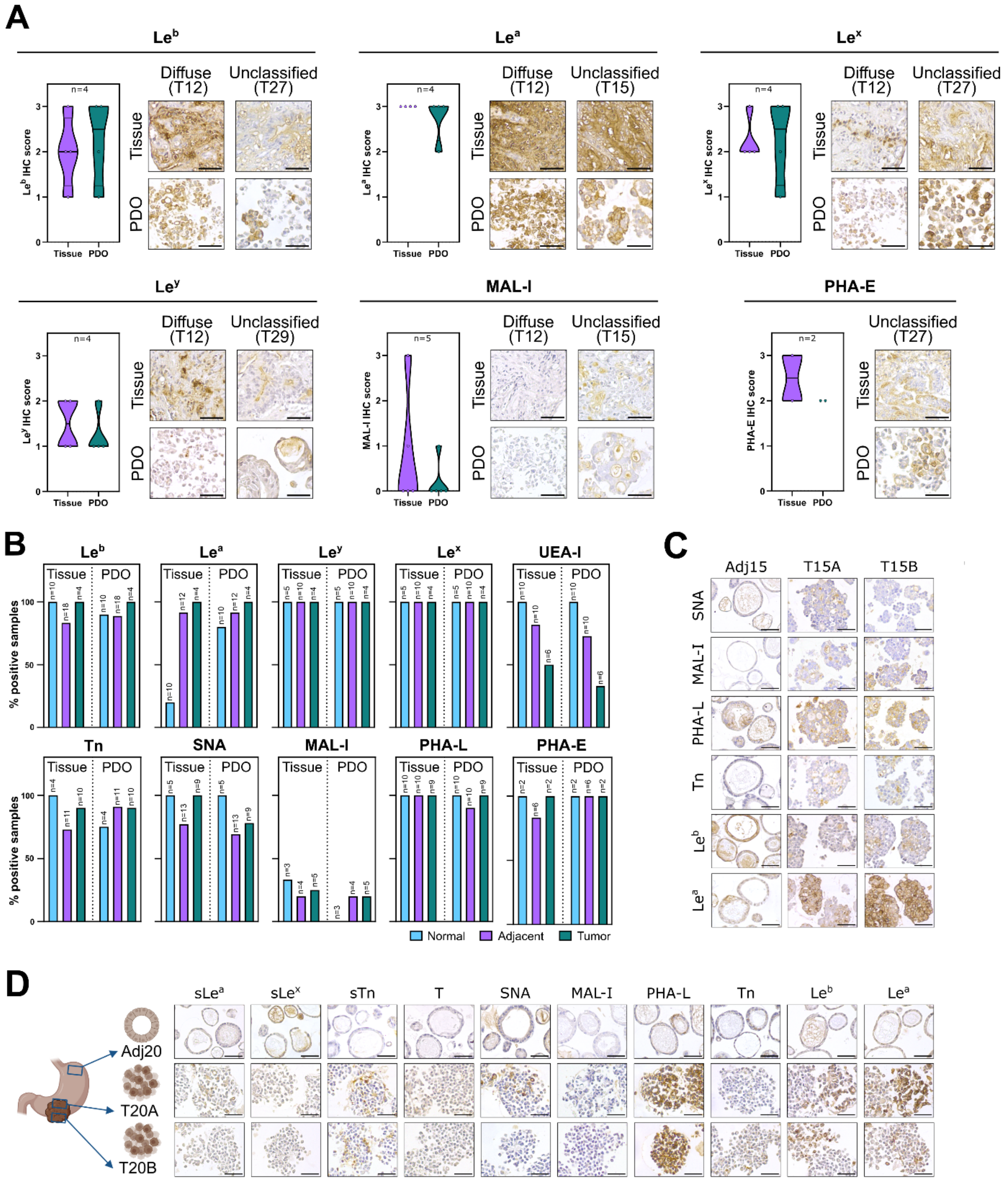

**Supplementary Figure 7. Tumor PDOs as *avatars* of the glycophenotype heterogeneity of the parental tissues.** Glycan profile analysis of parental tissues and corresponding PDO of additional glycan epitopes: including type I Lewis ($Le^a$, $Le^b$) and type II Lewis ($Le^x$, $Le^y$) antigens, α2,3 sialylated LacNAc structures (MAL-I), and bisecting *N*-glycans (PHA-E) structures. Scale bar corresponds to 50 µm. **B)** Percentage of normal, adjacent and tumor samples, including both tissues and receptive PDOs, showing positive staining for the glycosylation profile analyzed. **C)** Immunostaining for the represent glycan antigens of samples derived from the patient T15 (Adj15 and the two tumor siblings PDOs, T15A and

T15B). **D)** Immunostaining for the represent glycan antigens of samples derived from the patient T20 (Adj20 and the two tumor siblings PDOs, T20A and T20B). Scale bar corresponds to 50 µm

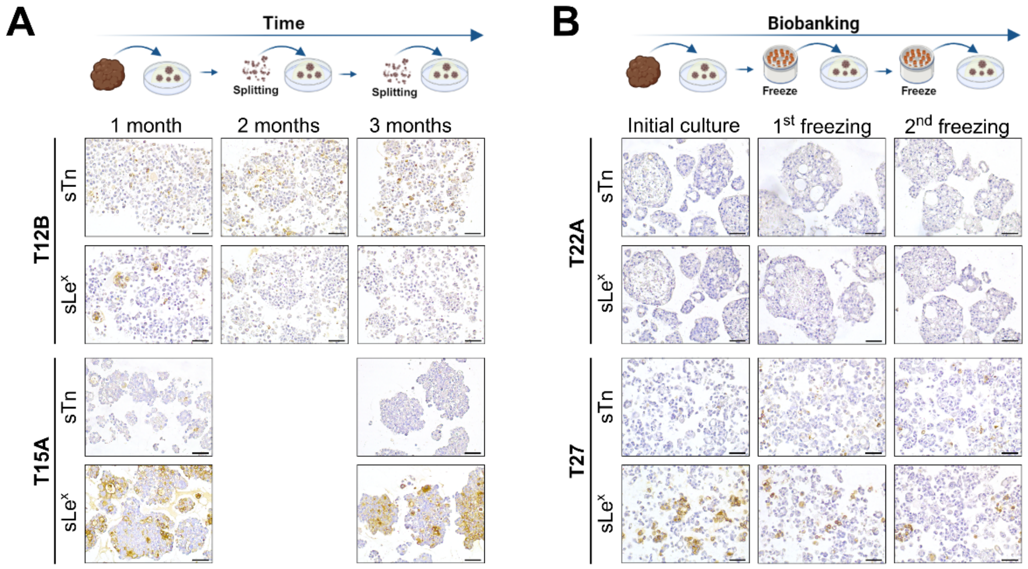


**Supplementary Figure 8. Tumor PDOs can maintain their glycan heterogeneity in culture over time and upon biobanking. A)** sTn and sLe$^x$ expression analysis over time in culture up to three months continuously**. B)** sTn and sLe$^x$ expression analysis upon biobanking and re-culture of PDOs. Scale bar corresponds to 50 μm.

### SUPPLEMENTARY DATA_SUPPLEMENTARY MATERIAL AND METHODS

#### ***Human tissue samples***

Fresh gastric mucosa, and tumor and the corresponding adjacent mucosa (0.2-1 $cm^3$ in size) were obtained from individuals undergoing sleeve resection or surgical gastrectomy/tumor resection at Centro Hospitalar e Universitário de São João (CHUSJ) in Porto, Portugal. Adjacent tumor mucosa tissue was collected approximately 1-2 cm away from the tumor tissue without macroscopic indication of any pathology. One-half of each patient's sample was fixed and embedded in paraffin for subsequent sectioning and histological analysis. The remaining half of the tissue was used for organoid culture. A total of 70 human tissue samples were used for this work. Among these samples, 11 gastric mucosa samples were obtained from 11 non-tumoral obese patient donors (N). Additionally, 29 samples were collected from gastric mucosa adjacent to the tumor (Adj) from 26 GC patients. The remaining 30 samples were obtained from tumor tissue (T) from 25 patients diagnosed with the following Lauren GC classification: intestinal subtype (44% of the patients), diffuse subtype (8% of the patients), and unclassified subtype (48% of the patients). All sample collection procedures were conducted upon approval by the CHUSJ ethical committee under reference number CES 223/2021 and following the patients' informed consent. An overview of the clinical data associated with the patient cohort used for the PDO biobank establishment is provided in **Supplementary Table 1** and **Supplementary Table 2**.

#### ***Gcnt1 knockout mice models***

The *Gcnt1*$^{-/-}$ mice strains were obtained from the Consortium for Functional Glycomics (CFG)[1]. Mice were bred and housed at the i3S animal house. Food and water were provided *ad libitum*. All animal experiments were performed in accordance with the European guidelines for the Care and Use of Laboratory Animals (directive 2010/63/UE), following a protocol approved by the i3S Animal Ethics Committee and the Portuguese National Authority for Animal Health (reference 2021_05). C57BL/6 *wild-type* and *Gcnt1*$^{-/-}$ (6–8-week-old) mice were euthanized by $CO_2$ inhalation and stomachs were harvested, washed with a 0.9 % NaCl solution and divided: half of the stomach collected in PBS for organoid generation and the other half fixed for immunostaining analysis.

***Generation of gastric mice and human non-tumoral patient-derived organoids***

Patient-derived organoids (PDOs) derived from normal human gastric mucosa and from mucosa adjacent to the tumor were generated as previously described[2]. Briefly, a small fragment of fresh gastric mucosa or tumor adjacent mucosa was washed with cold chelating buffer (5.6 mmol/L $Na_2HPO_4$, 8.0 mmol/L $KH_2PO_4$, 96.2 mmol/L NaCl, 1.6 mmol/L KCl, 43.4 mmol/L sucrose, 54.9 mmol/L D-sorbitol, 0.5 mmol/L DL-dithiothreitol, with a pH 7.0). The tissue fragment, free of the mucous and muscle layer, was cut into small pieces and washed vigorously in 10 mL of chelating buffer (5-10 times). Then, the tissue fragments were incubated in 20 mL of chelating buffer containing 10 mM EDTA for 5 minutes for mice tissues or 15 minutes for human tissues at room temperature (RT). After glands were released from the gastric tissue by squeezing the fragmented tissue using a glass microscopy slide, both glands and tissue pieces were collected in 30 mL of cold Advanced Dulbecco's modified Eagle (DMEM)/F12 medium (D8437, Sigma-Aldrich). The large tissue fragments were allowed to settle, and the cloudy supernatant containing glands was centrifuged at 200 g and 4 ºC for 5 minutes. Isolated gastric glands and fragments of gastric epithelium were mixed with ice-cold Matrigel (356231, Corning®) and seeded in pre-warmed 24-well plates. The Matrigel-cell mixture dome was left at 37 ºC to solidify for 10-30 minutes. Mice-derived organoids (MDOs) with differential core 1 glycan extension were generated from the stomach of C57BL/6 *wild-type* and *Gcnt1*$^{-/-}$ mice strain, as described above. A final volume of 500 µl of gastric organoid medium containing essential growth factors [DMEM/F12, 50 % WNT3A conditioned medium, 10 % of Noggin conditioned medium, 10 % of R-spondin1 conditioned medium, FGF10 200 ng/mL (78037.1, Stemcell™), Primocin 100 µg/mL (ant-pm-1, Invivogen), B-27™ supplement 1 X (17504-044, Thermo Fisher Scientific), EGF 50 ng/mL (PHG6045, Gibco™), Gastrin I 1 nM (Tocris), TGFβi 2 µM (A-83-01, SML0788, Sigma-Aldrich), N-acetyl-L-cystein 1 mM (NAC, A9165, Sigma-Aldrich), RHOKi 10 µM (Y-27632, ab120129, Abcam)] was added to each well. The conditioned media used in this work were produced using in-house cell lines that stably express Wnt3A, R-spondin1, and Noggin kindly provided by Dr Bartfeld lab[3]. The organoids were left to grow at 37 ºC and 5 % of $CO^2$ in a humidified incubator. Fresh medium was added every 2-3 days and organoids were passaged every 7-12 days at a ratio of 1:4 to 1:8.

***Generation of tumor patient-derived organoids***

The protocol for the generation of tumor PDOs was adapted from Bartfeld *et al.* (2015)[2] and Wallaschek *et al.* (2019)[3]. Firstly, a small fragment of tumor tissue (avoiding normal mucosa contamination) was washed with ice-cold PBS and minced using scissors and blades. Then, the tumor fragments were digested at 37 °C in agitation in a digestion buffer containing advanced DMEM/F12 (D8437, Sigma-Aldrich) with 1.5 mg/mL of collagenase A (10103578001, Roche), 20 µg/mL of hyaluronidase (H4272, Sigma-Aldrich) and 100 µg/mL of primocin (ant-pm-1, Invivogen). The incubation time differed between samples from 30 minutes to 2 hours depending on the tissue stiffness. The tumor digested fragments were then disrupted mechanically by up-and-down using a 1000 µL pipette and incubated with Tryple Express (12605028, Gibco™) for 15 minutes at 37 °C for cell dissociation. Cell clusters and remaining fragments were washed using DMEM/F12 medium by centrifugation at 1500 rpm for 5 minutes and resuspended in 40 µL of Matrigel (356231, Corning®) per well. Whenever there was evidence of possible normal gastric organoid growth, WNT3A was removed from the culture medium for at least 4 weeks. Fresh medium was added every 2-3 days and organoids were passaged every 10-15 days at a ratio of 1:2 to 1:8 depending on the organoid line. Additionally, two organoid lines were derived from patient-derived xenografts (PDX).

***Generation of patient-derived tumor organoids xenografts***

To develop gastric patient-derived tumor organoids xenografts (PDOXs), a surgical subcutaneous dorsal implantation was performed as described previously[4]. Briefly, 50 µL suspension of a given organoid dissolved in a Matrigel matrix/PBS mixture (~10.000 PDOs) was injected in nude CBA mice (i3S animal facility, Portugal). Eleven tumor-PDOs were used with different histologic phenotypes: T1, T1.1 – solid pattern; T10, T22A, T22B, T23, – glandular pattern, T15B– solid and glandular pattern, T12– non-cohesive pattern, and T20A, T20B - non-cohesive with solid pattern. See **Supplementary Table 2** for clinical-pathologic data. The study was performed and approved by the Animal-based studies Ethical Committee and fulfilled all legal requirements (i3S Animal Ethical Committee and DGAV, Portugal).

***Confocal and phenotypic analysis***

Free-floating fixed organoids were permeabilized with 50 mM of $NH_4Cl$ for 15 minutes and with PBS 0.3 % Triton X-100 (T9284, Sigma Aldrich) for 15 minutes in rotation followed by washing steps using PBS 0.1 % Triton X-100. Samples were blocked with the UltraVision Protein Block (TA125PBQ, Thermo Scientific) for 15 minutes. For cytoskeletal (actin filaments) staining organoids were incubated with CruzFluor™ 594 phalloidin conjugate (sc-363795, Santa Cruz Biotechnology) for 30 minutes diluted

1:1000 and the nuclei were stained with PhenoVue Hoechst 33342 (PELSCP71, Perkin Elmer) for 30 minutes diluted 1:1500 at RT. Finally, organoids suspensions were visualized using the Opera Phenix Plus (Perkin Elmer). Images were acquired using the Harmony High-Content Imaging and Analysis Software (Perkin Elmer).

***Type I and type II Lewis modulation in non-tumoral patient-derived organoids***

Glycan expression modulation/differentiation analysis was performed in normal human organoids with 7 days of grown passaged at a ratio of 1:6. Non-tumoral organoids (N2, N11, Adj9) were incubated with Tryple Express (12605028, Gibco™) for 15 minutes at 37 °C for a complete cell dissociation and embedded in Matrigel. To mimic the differentiation observed in gastric mucosa, we differentiated organoids into two different types according to the cell's expression of the epithelium cell lineages (MUC5AC and MUC6-expressing organoids). To differentiate organoids into Lewis type I (MUC5AC-positive) or into Lewis type II (MUC6-positive) the WNT3A concentration in the organoid medium was altered. For that, organoids were allowed to grow for 10 days in the presence of 50 % of WNT3A to differentiate organoids into mixed-type (MUC5AC and MUC6-positive), while the foveolar pit-type organoid differentiation (MUC5AC-positive) was induced by removing WNT3A from the culture medium at day 4 of the development and cultures were kept for 6 days under these conditions before collection. Undifferentiated organoids were collected at day 4 after plating.

***Helicobacter pylori strains and binding assay***

The *H. pylori* strains 17875/$Le^b$ and 17875babA1::kan babA2::cam (17875babA1A2) were obtained from the Department of Medical Biochemistry and Biophysics, Umeå University, Sweden[5]. The 17875/$Le^b$ strain is a spontaneous mutant that binds $Le^b$ but does not bind to sialylated antigens[5], while *double mutant* strain 17875babA1::kan babA2::cam (17875 DM.) binds to sialylated structures as the case of $sLe^x$. *H. pylori* strains were labeled with FITC, and adhesion assays were performed as previously described[6, 7]. Briefly, sections were rehydrated followed by incubation for 1 hour with blocking buffer: 1 % BSA in PBS containing 0.05 % Tween 20. The FITC-labeled bacterial suspension was diluted 5-fold in the blocking buffer and 100 µL was used to incubate each sample for 2 hours at RT. Slides were subsequently washed three times with PBS containing 0.05 % Tween 20 and nuclear counterstaining was performed with DAPI. The images were acquired using a Zeiss Axio cam MRm and the AxioVision Rel.

4.8 software (both from Zeiss). Evaluation of bacterial binding in both tissues and PDOs was assessed by visual observation and using Fiji software[8]. For the PDO analysis, ~12 organoids per sample were used to quantify bacterial binding in the epithelial cell layer. Selection of regions of interest (ROI) and the measurement of the area ($\mu m^2$) of those regions was performed using the following: Analyze -> Tools ->ROI Manager->Polygon selections-> ROI Manager (Measure). For this study, each PDO was selected as one ROI. The number of foci (*H. pylori*) presented in each PDO/ROI was assessed using the following: Process-> Find Maxima ->Prominence >1800. The number of foci was divided by the area of each ROI (foci/ $\mu m^2$).

***RNA organoid extraction, cDNA synthesis and quantitative PCR***

For PDOs RNA extraction, organoids were released from Matrigel using a Corning® Cell Recovery Solution (354253, Corning®) for 45 at 4°C, followed by washing with PBS twice and pelleted. Total RNA from organoids pellets was purified using the PureLink RNA Mini Kit (ThermoFisher) following manufactures' instructions. The eluted RNA was quantified and qualified using the NanoDrop™ One/OneC Microvolume UV-Vis Spectrophotometer from Thermo Fisher Scientific. The complementary DNA (cDNA) was obtained from reverse transcription using the SuperScript IV First-Strand cDNA Synthesis Reaction, using a total of 500 ng of the extracted RNA. Then, the samples were subjected to RT-qPCR amplification using the iTaq™ Universal SYBR® Green Supermix (Bio-Rad). The amplification process was performed on the real-time cycler CFX96TM (Bio-Rad). Each reaction mixture included the following components: 6 µL of DNAse/RNAse free treated $H_2O$, 1 µL of the respective forward and reverse primer (10 µM) (listed in **Supplementary Table 3**) and 10 µL of SYBR Green. Initially, 2 µL of cDNA diluted 1:2 was added to each well of a 96-well PCR plate, followed by the addition of the corresponding mixture, resulting in a total volume of 20 µL per reaction. PCR products were then run in a 2 % agarose gel. β-actin (*ACTB*) was used as a reference housekeeping gene.

***Immunohistochemistry, immunofluorescence and lectin staining***

For histological analysis, organoids were released from Matrigel using Corning® Cell Recovery Solution (354253, Corning®) at 4 °C for 45 minutes, followed by three washes with PBS followed by fixation using 4 % paraformaldehyde (PFA, Alfa Aesar) solution overnight at 4 °C. For immunohistochemistry analysis fixed organoids were included in histogel before paraffin embedding. Tissues were fixed in 10 % buffered formalin and embedded in paraffin.

Paraffin-embedded sections of the patient tissues and PDOs were stained with hematoxylin and eosin (H&E) for histological examination. Briefly, paraffin sections were deparaffinized and rehydrated, followed by heat-induced antigen retrieval with citrate buffer (10 mM, pH = 6.0) when required or with EDTA buffer (pH = 8.0) for CDX2 staining, and neutralization of endogenous peroxidase activity with 3 % $H_2O_2$ in methanol for the immunohistochemistry analysis. To block non-specific background staining, slides were incubated for 30 minutes at RT with UltraVision Protein Block (TA-125-PBQ, Thermo Fisher Scientific) for antibody staining or with 10 % BSA for lectin staining. Tissue sections were incubated overnight at 4 ºC with primary antibodies or 60 minutes at RT with biotinylated lectins. The glycoprofile characterization included type I Lewis (Lewis a ($Le^a$), Lewis b ($Le^b$), sialyl Lewis a ($sLe^a$)) and type II Lewis (Lewis x ($Le^x$), Lewis y ($Le^y$), $sLe^x$) antigens, fucosylated (*Ulex europaeus agglutinin-I* - UEA-I: type 2 blood group H, Fucα1,2Galβ1,4GlcNAc), sialylated structures (*Sambucus nigra* agglutinin- SNA: α2,6-sialylated Lac(di)NAc; *Maackia amurensis* lectin I - MAL-I: α2,3 sialylated LacNAc), branched *N*-glycans (*Phaseolus vulgaris* leucoagglutinin - PHA-L: β1,6 branched *N*-glycans), bisecting *N*-glycans (*Phaseolus vulgaris* erythroagglutinin - PHA-E: bisecting GlcNAc with type 2 LacNAc), and short truncated *O*-glycans (T, Tn, and sTn) structures. Specific antibodies and lectins used are described in **Supplementary Table 4.**

For immunohistochemistry analysis, slides were incubated for 30 minutes with the respective biotin-labeled secondary antibody (**Supplementary Table 4**) and incubated with Vectastain® ABC reagents (Vector Laboratories) for an additional 30 minutes at RT. For lectin staining, slides were directly incubated with Vectastain® ABC reagents for 30 minutes at RT. Slides underwent chromogenic staining with DAB containing 0.01 % $H_2O_2$ and nuclear staining was performed using Mayers' hematoxylin solution. The CDX2 immunostaining was performed in an automated Ventana BenchMark ULTRAStaining System, using OptiView DAB IHC Detection Kit (Roche/Ventana Medical Systems), according to the manufacturer's instructions. Digital pictures were taken under 200X magnification with Leica DM2000 LED (Leica Microsystems).

For immunofluorescence analysis, after primary antibody incubation, slides were incubated for 60 minutes in the dark with respective Alexa Fluor® conjugated secondary antibodies diluted 1:500 in PBS 5 % BSA (**Supplementary Table 4**). Slides were mounted in VectaShield mounting medium containing DAPI for nuclear staining (Vector Laboratories). Images were acquired using a Zeiss Axio cam MRm and the AxioVision Rel. 4.8 software (both from Zeiss).
